# Super-enhancers require a combination of classical enhancers and novel facilitator elements to drive high levels of gene expression

**DOI:** 10.1101/2022.06.20.496856

**Authors:** Joseph Blayney, Helena Francis, Brendan Camellato, Leslie Mitchell, Rosa Stolper, Jef Boeke, Douglas Higgs, Mira Kassouf

## Abstract

Super-enhancers (SEs) are a class of compound regulatory elements which control expression of key cell-identity genes. It remains unclear whether they are simply clusters of independent classical enhancers or whether SEs manifest emergent properties and should therefore be considered as a distinct class of element. Here, using synthetic biology and genome editing, we engineered the well characterised erythroid α-globin SE at the endogenous α-globin locus, removing all SE constituent elements in a mouse embryonic stem cell-line, to create a “blank canvas”. This has allowed us to re-build the SE through individual and combinatorial reinsertion of its five elements (R1, R2, R3, Rm, R4), to test the importance of each constituent’s sequence and position within the locus. Each re-inserted element independently creates a region of open chromatin and binds its normal repertoire of transcription factors; however, we found a high degree of functional interdependence between the five constituents. Surprisingly, the two strongest α-globin enhancers (R1 and R2) act sub-optimally both on their own and in combination, and although the other three elements (R3, Rm and R4) exhibit no discernible enhancer activity, they each exert a major positive effect in facilitating the activity of the classical enhancers (R1 and R2). This effect depends not simply on the sequence of each element but on their positions within the cluster. We propose that these “facilitators” are a novel form of regulatory element, important for ensuring the full activity of SEs, but distinct from conventional enhancer elements.

## Introduction

Super-enhancers (SEs) are dense clusters of regulatory elements with the bioinformatic signatures of enhancers; they recruit unusually high levels of co-activators and associated chromatin modifications, and regulate genes lying 10s-1000s kb away in the genome (Whyte et al., 2013). SEs are often regulators of cell identity genes and are frequently mutated in association with complex traits and genetic diseases (Dębek & Juszczyński, 2022; Harteveld & Higgs, 2010; Higgs et al., 2012; Tang et al., 2020; Yamagata et al., 2020). Despite extensive analysis, it remains unclear whether SEs are merely groups of classical enhancers, or whether they contain both enhancers and other types of functionally distinct regulatory elements, together cooperating to up-regulate their target gene(s) (Blobel et al., 2021; Grosveld et al., 2021; Moorthy et al., 2017; Pott & Lieb, 2015; Wang et al., 2019).

Detailed analysis of SEs is challenging since many drive expression of genes playing central roles in complex transcriptional and epigenetic programmes. Consequently, analysing changes in SE-regulated gene expression upon SE engineering is often confounded by associated changes in cell lineage and differentiation. In addition, previous studies analysing SEs have drawn conclusions from the deletion of just one or two constituent elements and many studies have relied on artificial reporter-based assays divorced from their functionally relevant native chromatin contexts. To date, a number of studies have dissected SEs in detail (Bender et al., 2012; Grosveld et al., 2021; Hay et al., 2016; Hnisz et al., 2015; Hörnblad et al., 2021; Huang et al., 2018; Shin et al., 2016; Thomas et al., 2021), but further, more rigorous analysis of model SEs is essential to determine their true nature.

Here, we present a series of genetic models rebuilding a SE from an enhancerless baseline and show that this degree of comprehensive dissection is essential to fully determine the complex interdependencies between each constituent element. To do this, we required a well characterized, tractable SE, in which phenotypes arising from extensive genetic engineering could be easily interpreted. The mouse α-globin SE (α-SE) provides an ideal model; it lies together with the duplicated α-globin genes in a well-defined 65kb sub-TAD and up-regulates α-globin gene expression during erythropoiesis. The α-SE is exclusively activated during terminal erythroid differentiation, and its modification has no effect on erythroid cell identity or differentiation (Hay et al., 2016). Therefore, unlike most SE driven loci that have been analysed, all changes in gene expression can be directly related to the engineered changes in the SE alone (Oudelaar et al., 2021). We previously dissected the α-SE by individual and selective pairwise deletion of its five constituent elements: R1, R2, R3, R4, and Rm (a mouse-specific element). Importantly, in these experiments, elements were removed from an otherwise intact SE, and using this approach they appeared to act in an additive manner (**fig 1, A**). Enhancers R1 and R2 were identified as the two major activators of α-globin transcription contributing 40% and 50% to the SE’s total activity respectively. Despite showing conservation in sequence and synteny over ∼70 million years of evolution (Hughes et al., 2005), R3 and R4 display little or no inherent enhancer activity when removed individually from the endogenous locus or in classical enhancer reporter assays, and the same was true of Rm (Hay et al., 2016). We concluded that a more rigorous test of each element’s role would be to assay their abilities to up-regulate their cognate target gene, in their native chromosomal environments and developmental contexts after all other elements have been removed from the locus. If a SE constituent functions independently, its intrinsic ability to activate gene expression should match the change in expression following its deletion. For example, removal of R2 from the SE reduces α-globin expression by 50% (Hay et al., 2016); if it acts as an independent additive enhancer, a locus containing R2 alone would be expected to drive α-globin expression at 50% of its normal level (**fig 1, B**).

**Figure 1.**
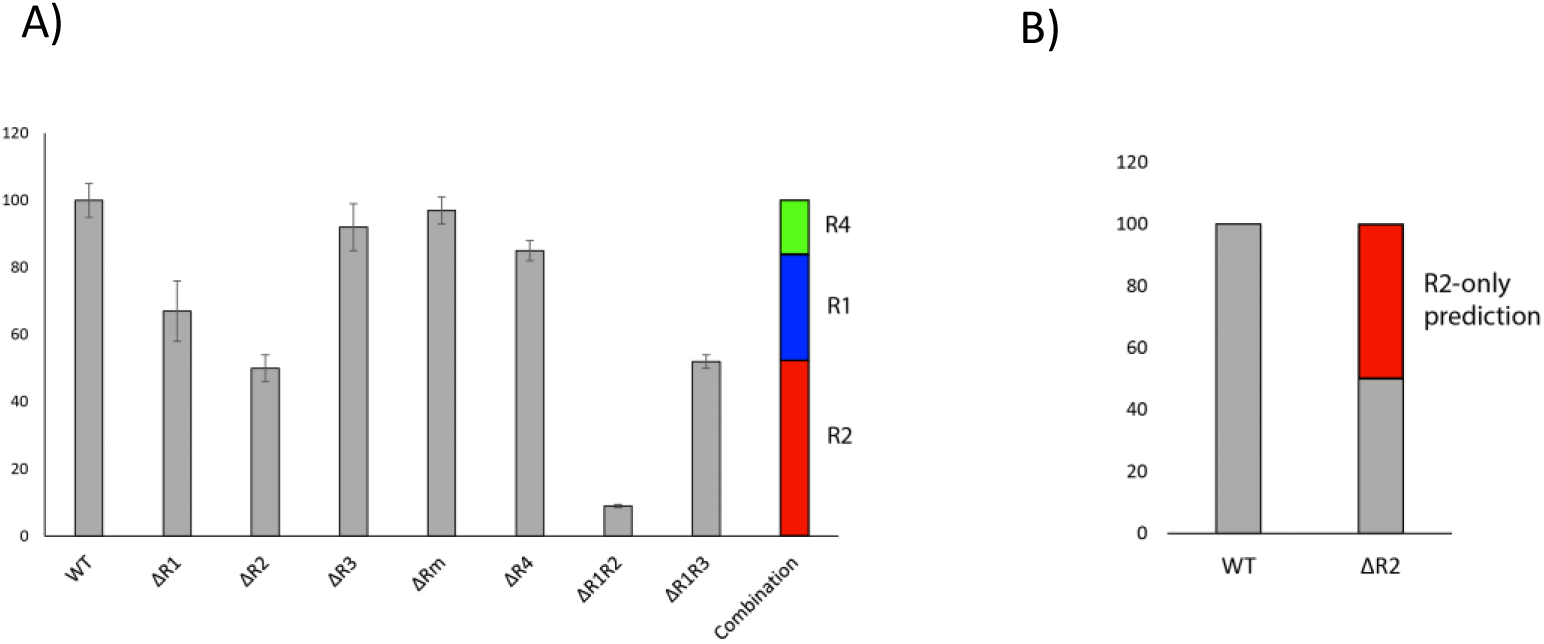
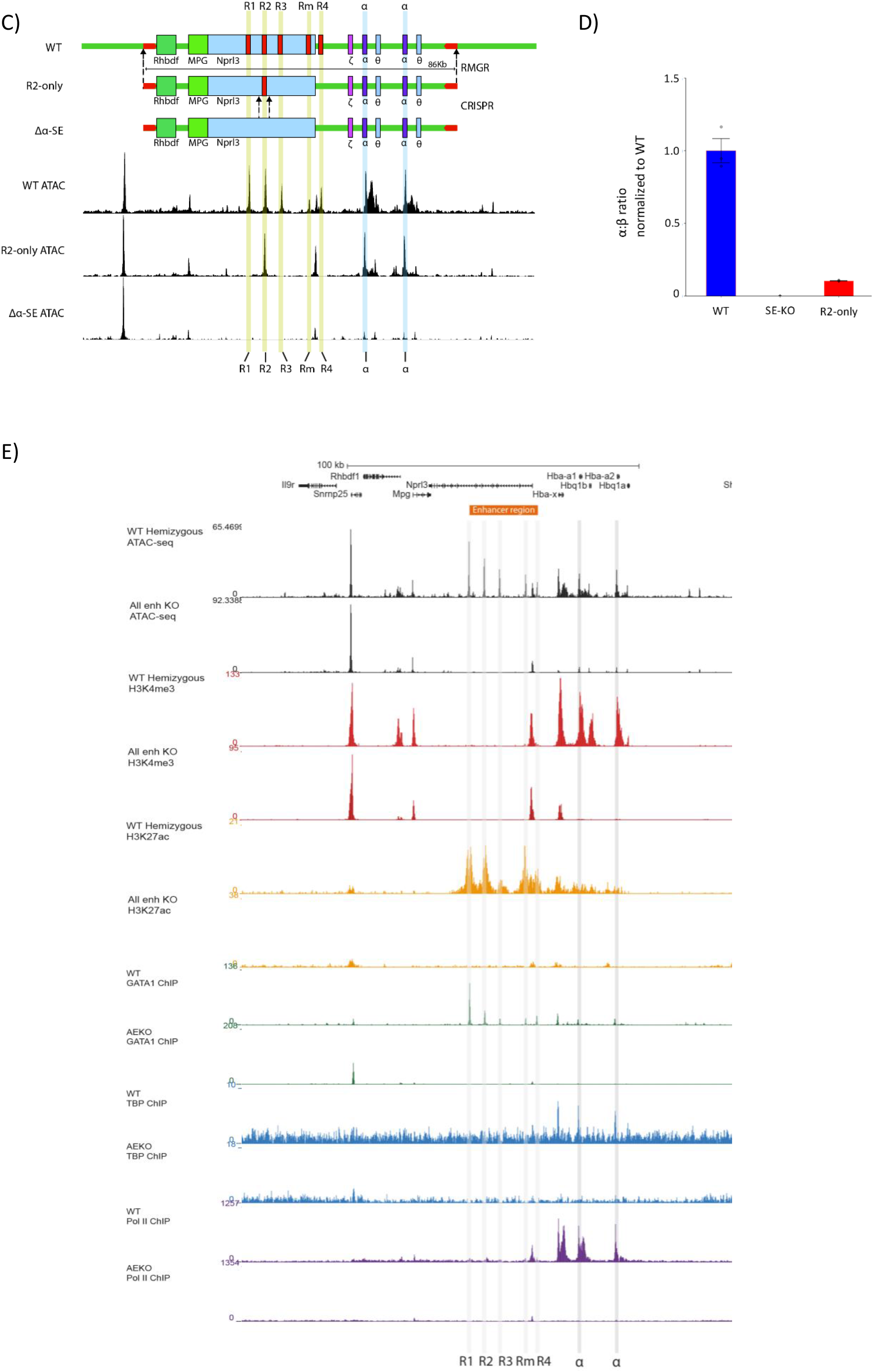
Generation of an R2-only mouse model to test the sufficiency of the R2 enhancer element. A) α-globin gene expression from seven mouse models harboring homozygous single or double enhancer element deletions (adapted from Hay et al., 2016). Far right: contribution of each enhancer element, calculated by subtracting each deletion model from WT. B) Prediction of R2-only α-globin gene expression, calculated by subtracting ΔR2 α-globin expression from WT. C) Design of the R2-only and Δα-SE α-globin loci. R2-only locus synthesized, assembled into a bacterial artificial chromosome, and delivered into the WT α-globin locus through recombination-mediated genomic replacement. R2 enhancer later deleted using CRISPR. Top = schematic of R2 - only RMGR, followed by R2 deletion; bottom = WT, R2-only and Δα-SE ATAC-seq. D) α-globin gene expression in EB-derived WT, Δα-SE and R2-only erythroid cells (n≥3) assayed by RT-qPCR. Expression normalized to β-globin and displayed as a proportion of WT expression. Dots = biological replicates; error bars = SE. E) ATAC-seq in EB-derived WT, Δα-SE in EB-derived erythroid cells (top), ChIPmentation for H3K4Me3, H3K27Ac, Gata1, TBP and Pol2 (beneath) (n=3).

To test this hypothesis, we engineered the endogenous α-globin locus in mouse embryonic stem (ES) cells to delete all five α-SE constituents (Δα-SE). This provides a “blank canvas” into which any combination of elements, in any position, can be re-introduced and the effects on α-globin expression assessed. Contrary to expectation, in *in vitro* differentiated erythroid cells (Francis et al., 2022), reinsertion of the strongest constituent of the α-SE, R2 (R2-only), resulted in five-fold lower levels of transcription than predicted in the absence of R1, R3, Rm and R4. To investigate this further, we generated an R2-only mouse model. R2-only mice also expressed very little α-globin and were not viable. Upon separation from the R1, R3, Rm and R4 elements, R2 retains the epigenetic signature and tissue-specific TF recruitment of an active enhancer, but exhibits attenuated coactivator recruitment, enhancer-promoter interactions, and eRNA transcription. Rebuilding the α-SE from the enhancerless Δα-SE baseline revealed that both major activators (R1 and R2), individually and when combined, are insufficient to drive high levels of gene expression. Although R3, Rm and R4 display the bioinformatic hallmarks of enhancers (H3K27Ac, H3K4Me1, tissue-specific TF recruitment), they are incapable of driving α-globin expression upon reinsertion into the Δα-SE locus. Nevertheless, adding one or more of these elements to R1 and R2 considerably increased α-globin expression and ultimately restored full levels of expression. Importantly, the effect of these elements which play a critical role in the activity of a SE, was dependent on their position within the locus rather than their sequences. We propose that “facilitators”, such as R3, Rm and R4, are a novel form of regulatory element important for attaining the full activity of SEs.

### Engineering an enhancerless mouse α-globin cluster as a test bed for elements of the SE

We first aimed to establish mouse ES cells in which all elements of the SE were deleted using the precise coordinates described in previous deletion models (Hay et al., 2016). Engineering several independent mutations in a single allele using conventional editing is slow and complicated, requiring multiple steps to ensure that all mutations are precise and present in *cis* to one another. Multiple editing steps in a single cell line can also introduce “off target” effects which may compromise the ability of the model ES cell to divide and differentiate normally into erythroid cells or to generate a subsequent mouse model.

To overcome these issues, we used a recently developed protocol for *de novo* assembly of large DNA fragments (**supplementary fig 1, A**) (Mitchell et al., 2021) to design and synthesise 86kb alleles containing either the wild type (WT) α-globin sub-TAD or an equivalent allele in which only R2 remained with R1, R3, Rm and R4 deleted (R2-only). These two synthetic alleles were each integrated using recombinase mediated genomic replacement (RMGR) (Wallace et al., 2007) into ES cells in which one copy of the α-globin locus had been already deleted (**fig 1, C**). These hemizygous ES cells allow genomics analysis to be conducted specifically on each engineered locus as well as allowing more efficient genome editing. A third genetic model was made by deleting the remaining R2 element from R2-only ES cells using a CRISPR-Cas9 approach, creating a model in which all elements of the α-globin SE had been removed from the locus (Δα−SE) (**fig 1, C**). We then used an embryoid body (EB)-based *in vitro* differentiation and erythroid purification system recently developed in our lab (Francis et al., 2022) to produce hemizygous WT, Δα−SE and R2-only erythroid cells in which the single remaining α-globin locus is derived from a synthetic construct.

Upon deletion of all five elements of the α-SE, EB-derived erythroid cells display an almost complete loss of α-globin expression (>99.9% loss), and all chromatin marks normally associated with the SE elements are absent (**fig 1, D, E**). Very small ATAC-seq peaks persist over the α-globin promoters but they are no longer bound by Gata1, Pol II or TBP, or marked by H3K4me3 (**fig 1, E**). In the absence of the enhancers, H3K27ac is almost entirely lost from the locus, with only a very small peak associated with the embryonic ζ-globin gene remaining. In summary, the Δα-SE model provides a well characterised baseline for studying the role of the SE elements individually and in combination during erythropoiesis.

### A single enhancer driven α-globin locus (R2-only) is associated with severe downregulation of α-globin expression and embryonic lethality

To determine the individual contribution of the R2 enhancer element to α-globin expression, we compared the structure and function of the enhancerless locus (Δα-SE) with a locus in which the R2 element is reinserted into Δα-SE ES cells, lying at its normal position in the absence of R1, R3, Rm and R4. In this case, all enhancer activity comes from R2 alone (R2-only) (**fig 1, C**). Previously, we showed that deleting R2 from the α-SE (ΔR2) causes a 50% reduction in α-globin transcription compared to WT (Hay et al., 2016). We therefore predicted that R2-only erythroid cells would produce 50% α−globin expression (**fig 1, B**). Unexpectedly, EB-derived R2-only cells expressed only 10% α-globin, five-fold less than predicted (**fig1, D**).

To investigate the R2-only phenotype further, we generated an R2-only mouse model, in which the endogenous α-SE is replaced with a SE containing R2 but not R1, R3, Rm or R4. Previous ΔR2 mice displayed a 50% reduction in α-globin expression and no significant changes in red cell parameters (Hay et al., 2016); we therefore originally expected R2-only mice to present with a similar gene expression and haematological phenotype. In contrast to ΔR2 mice, R2-only mice were largely non-viable. In 16 heterozygote crosses harvested at embryonic days E9.5, E10.5, E12.5, E14.5 and E17.5 the Mendelian ratio of wt:hets:homs were as expected (**supplementary table 1**); however, homozygous R2-only embryos were visibly smaller and paler than their WT and heterozygous littermates (**fig 2, A**). We obtained only one surviving homozygote with anaemia and severe splenomegaly, which died prematurely at 7 weeks.

**Figure 2.**
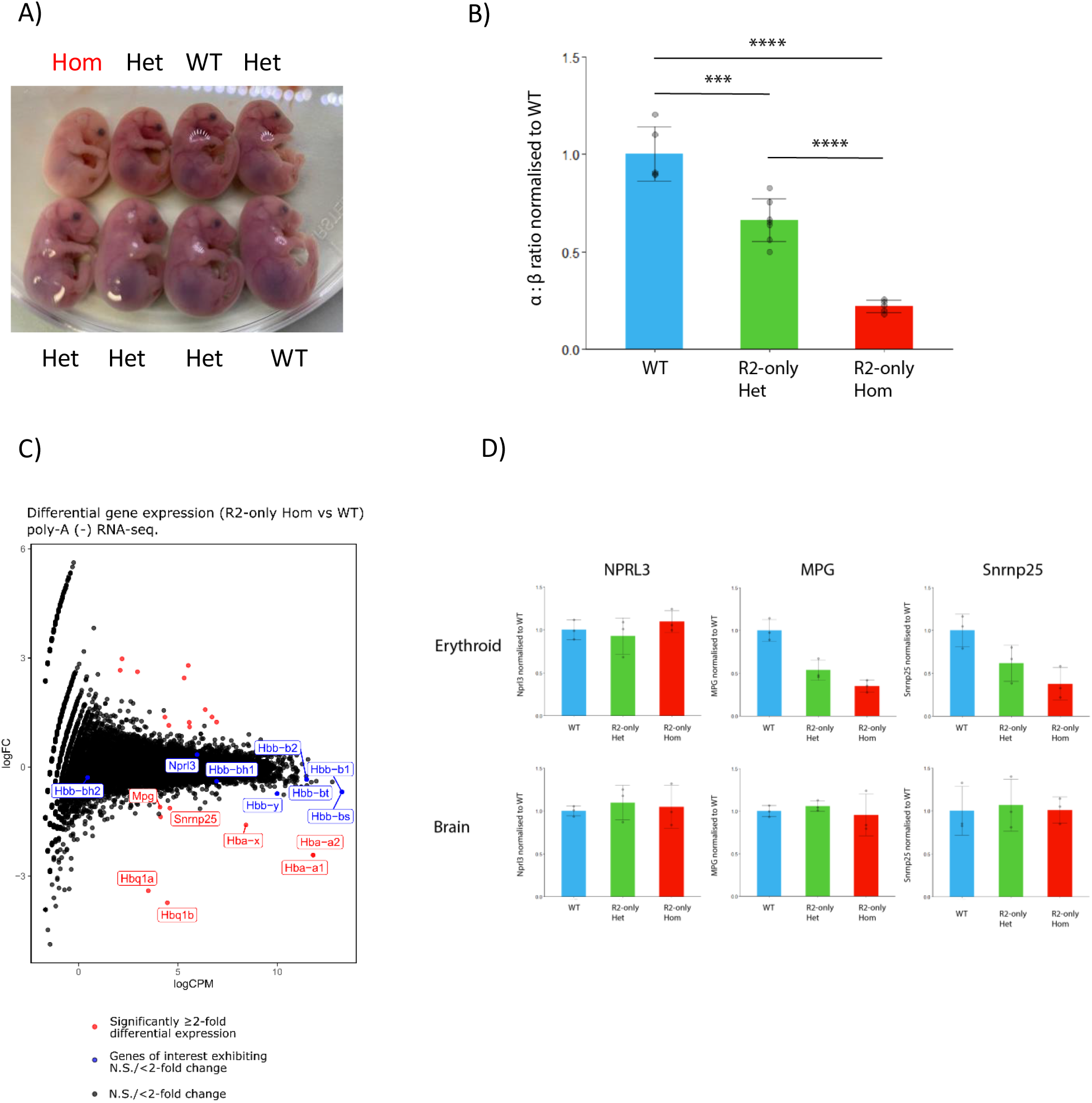
R2-only mice are inviable and exhibit severely attenuated α-globin expression. A) Representative image of R2-only Het X Het litter. Pregnant female sacrificed at embryonic day E17.5, and fetuses extracted. B) RT-qPCR comparing α-globin expression in foetal liver erythroid cells from WT, R2-only heterozygous and R2-only homozygous E12.5 littermates. Expression normalized to β-globin and displayed as a proportion of WT expression. Dots = biological replicates; error bars = SE. C) Poly-A minus RNA-sequencing comparing gene expression in foetal liver erythroid cells from WT (n=2) and R2-only homozygous (n=3) littermates. RNA was rRNA depleted, and poly-A minus transcripts isolated by removing poly-A positive fraction prior to library preparation. Red = statistically differentially expressed, blue = informative erythroid genes (non-differentially expressed). As well reductions in transcription of α-like globins, expression of Snrnp25 and MPG (two genes upstream of the 5’ boundary of the α-globin sub-TAD) was reduced. D) RT-qPCR comparing Nprl3, MPG and Snrnp25 expression in WT, R2-only heterozygous and R2-only homozygous littermates. Expression normalized to RPS18 and displayed as a proportion of WT expression. Dots = biological replicates; error bars = SE. Upper = E17.5 erythroid cells; lower = matched E17.5 Brain tissue.

The R2-only phenotype, with severe anaemia in homozygotes, suggested that removing R1, R3, Rm and R4 had compromised α-globin expression more than expected. To assess α-globin transcription we isolated E12.5 embryos, extracted fetal livers (FLs) (the definitive erythroid compartment at this stage), and performed RT-qPCR. R2-only erythroid cells expressed only 15% α-globin compared to WT littermates rather than the predicted 50% (**fig 2, B**). A similar downregulation of α-globin expression was observed at all developmental stages from E9.5-E17.5 (**supplementary fig 2, A**). Poly-A minus RNA-seq on FL erythroid cells from three R2-only and two WT littermates confirmed the RT-qPCR results, and showed that expression of various erythroid and developmental markers were unaffected in R2-only FL erythroid cells (**fig 2, C**). Some minor changes in the expression of two genes flanking the α-globin locus were noted (**fig 2, C**).

### The R2 element retains its enhancer identity in the absence of other SE elements but fails to recruit co-activators

To determine if R2 remains an active enhancer after removing R1, R3, Rm and R4, we examined the accessibility and epigenetic status of the α-globin locus. ATAC-seq revealed that R2 and both α-globin promoters (α-globin promoters) remain accessible in R2-only FL erythroid cells, and that R1, R3, Rm and R4 are by far the most differentially accessible regions genome-wide (**Fig 3, A, B**). ChIPmentation experiments in the R2-only model, using antibodies against H3K4Me1, H3K4Me3 and H3K27Ac, showed that R2 and the α-globin promoters are marked by active enhancer- (H3K4me1 and H3K27Ac) and promoter- (H3K4Me3 and H3K27Ac) associated histone modifications, respectively, albeit to a lesser extent than in WT cells (**Fig 3, B**). Furthermore, enhancers recruit high levels of tissue-specific TFs. Erythroid-specific TFs (e.g. Gata1 and Nf-e2) occupied both R2 and the α-globin promoters to an equivalent degree in R2-only and WT FL erythroid cells (**Fig 3, C**). We conclude that in the absence of other SE elements, R2 retains its identity as an enhancer, recruiting transcription factors and creating a region of open chromatin.

**Figure 3.**
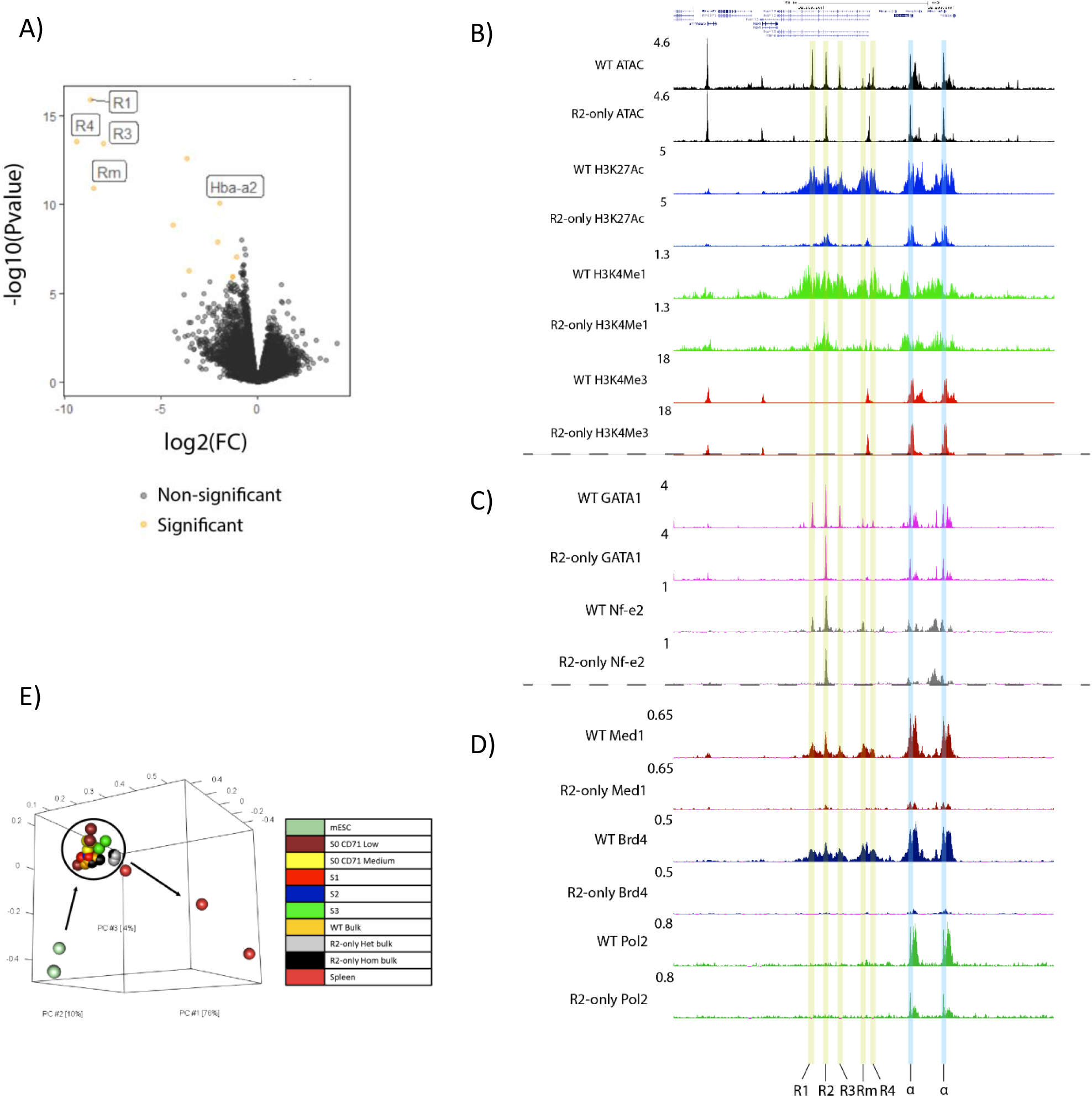
R2 retains the hallmarks of an active enhancer in R2-only erythroid cells, but coactivator recruitment is significantly reduced. A) Genome-wide differential accessibility assessed through ATAC-seq in foetal liver erythroid cells from WT (n=3) and R2-only homozygous (n=3) littermates. Yellow = significantly differentially accessible; black = non-significant. B) ATAC-seq in WT (n=3) and R2-only homozygous foetal liver erythroid cells (black); H3K27Ac ChIPmentation in WT (n=2) and R2-only homozygous (n=2) foetal liver erythroid cells (blue); H3K4Me1 ChIPmentation in WT (n=2) and R2-only homozygous (n=2) foetal liver erythroid cells (green); H3K4Me1 ChIPmentation in WT (n=2) and R2-only homozygous (n=2) foetal liver erythroid cells (red). Tracks = merged biological replicates. (Coordinates = chr11: 32,090,000-32,235,000). C) Gata1 ChIPmentation in WT (n=3) and R2-only homozygous (n=3) foetal liver erythroid cells (top); Nf-e2 ChIPmentation in WT (n=3) and R2-only homozygous (n=3) foetal liver erythroid cells (bottom). Tracks = merged biological replicates. (Coordinates = chr11: 32,090,000-32,235,000). D) Med1 ChIPmentation in WT (n=3) and R2-only homozygous (n=3) foetal liver erythroid cells (top); Brd4 ChIPmentation in WT (n=3) and R2-only homozygous (n=3) foetal liver erythroid cells (middle); Pol2 ChIPmentation in WT (n=2) and R2-only homozygous (n=2) foetal liver erythroid cells (bottom). Tracks = merged biological replicates. (Coordinates = chr11: 32,090,000-32,235,000). E) Principal component analysis comparing genome-wide ATAC-seq peaks in WT, R2-only homozygous and R2-only heterozygous foetal liver erythroid cells alongside mESC, developmentally-staged foetal liver erythroid cell, and adult spleen erythroid cell ATAC-seq peaks.

SEs are, in part, defined by the extent to which they recruit high levels of transcriptional coactivators (Whyte et al., 2013). To investigate R2’s capacity to recruit coactivators in the presence/absence of the other four α-SE constituents, we performed ChIPmentation with antibodies against Med1, a member of the mediator complex, and bromodomain-containing protein 4 (Brd4), a transcriptional and epigenetic regulator. WT FLs recruit high levels of Med1 and Brd4 to the α-SE and α-globin promoters, but in R2-only FL erythroid cells, recruitment of both factors was severely reduced (**Fig 3, D**). Mediator plays a central role in Pol2 recruitment and stability at the promoter; therefore, we asked whether reduced Med1 occupancy at the α-globin promoters corresponds with changes to formation of the preinitiation complex. We performed ChIPmentation experiments with antibodies against TATA-binding protein (TBP) and Pol2. There was no change in TBP recruitment in R2-only erythroid cells, consistent with its autonomous DNA-binding activity; however, there was a substantial reduction in Pol2 occupancy at both α-globin promoters (**supplementary fig 3, A**).

### R2 eRNA transcription is reduced in the absence of R1, R3, Rm and R4

Recent studies have discovered that enhancers are actively transcribed, producing bidirectional transcripts of varying lengths (Sartorelli & Lauberth, 2020). Enhancer transcription appears to be related to enhancer activity, although whether eRNAs have any function, or if the relationship between transcription and enhancer strength is merely correlative, remains unclear (Arnold et al., 2020). To explore eRNA transcription from the R2 element, we analysed the poly-A minus RNA-seq data. Because R2 is located in an intron of the Nprl3 gene, which is active in erythroid cells and transcribed on the negative strand, we had to restrict our investigation of R2 transcription to the positive strand. In WT FL cells, we found clear transcripts originating from all five α-SE constituents (**supplementary fig 4, A**), whereas in R2-only cells, only the R2 enhancer showed any evidence of transcription.

To compare R2 eRNA transcription in WT and R2-only cells quantitatively, we performed a “virtual qPCR”, normalizing levels of R2 eRNA to eRNA originating from the β-globin HS2 enhancer, a member of the β-globin locus control region (LCR) and a well characterised SE. This revealed a ∼3-fold reduction in R2 eRNA transcription in R2-only cells compared to WT (**fig 4, A**).

**Figure 4.**
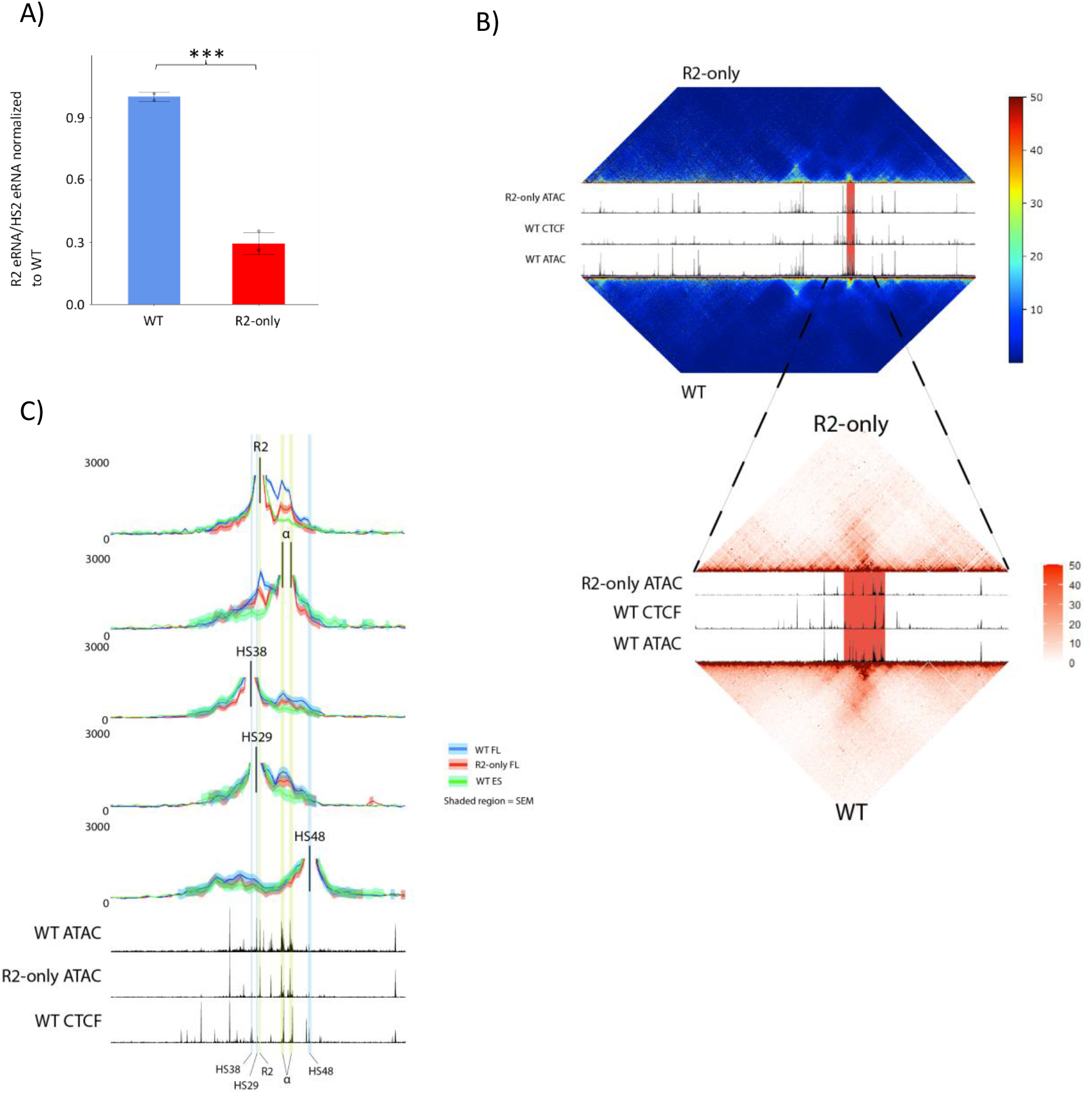
R2-α-globin promoter interaction frequency and R2 eRNA transcription are reduced in R2-only erythroid cells. A) Virtual qPCR conducted on poly-A minus RNA-seq, comparing R2 eRNA expression in WT (n=2) and R2-only homozygous (n=3) foetal liver erythroid cells (see methods). Transcripts originating from the R2 enhancer normalized to those originating from the HS2 enhancer within the β-globin LCR and displayed as a proportion of WT expression. Dots = biological replicates; error bars = SE. B) Tiled-C heatmaps comparing all-vs-all interaction frequency throughout the α-globin locus in R2-only homozygous (n=3) and WT (n=2) foetal liver erythroid cells. Upper: coordinates = chr11:29,900,000–33,230,000. Lower: zoomed heatmap over the α-globin locus; coordinates = chr11:31900000-32400000. Heatmaps = merged biological replicates. Bins = 2,000bp; tracks = R2-only homozygous ATAC-seq; WT CTCF ChIP-seq; WT ATAC-seq (top-bottom); red = α-globin locus. C) Virtual capture plots: pairwise interactions throughout the zoomed tiled locus (chr11:31900000-32400000) in which viewpoints participate. Viewpoints: R2 enhancer, α-globin promoters (considered together due to similarity in sequence, see methods), HS38 CTCF site, HS29 CTCF site, HS48 CTCF site (top-bottom). Blue = WT foetal liver erythroid cells (n=2), red = R2-only homozygous foetal liver erythroid cells (n=3), green = mESC cells (n=3).

### Enhancer-promoter interaction is compromised in the absence of the other α-SE constituents

Numerous publications have demonstrated high frequency interactions between SEs and their cognate target genes (Allahyar et al., 2018; Beagrie et al., 2017; Grosveld et al., 2021; Ing-Simmons et al., 2015; Li et al., 2018; Oudelaar & Higgs, 2021). These interactions appear crucial for effective up-regulation, of the target gene, although the mechanism(s) facilitating SE-target gene interaction, and the spatiotemporal relationship between interaction and activation, remain unclear. Indeed, previous chromatin conformation capture (3C)-based studies have shown that in erythroid cells the α-SE constituents, particularly R1 and R2, interact frequently with the α-globin promoters (Hanssen et al., 2017; Hay et al., 2016; Hua et al., 2021; Hughes et al., 2014; King et al., 2021; Oudelaar et al., 2018, 2019). To investigate whether R2’s ability to contact the α-globin promoters is affected in the R2-only locus in erythroid cells, we performed tiled-C: a low-input, high-resolution 3C-based technique, which allows comparison of “all-vs-all” pairwise chromatin interactions at a specific genomic locus (in this case, 3.3 Mb surrounding α-globin) (Oudelaar et al., 2020).

Visual inspection of chromatin interaction heat maps suggested a reduction in the overall frequency of pairwise interactions throughout the α-globin sub-TAD in R2-only FL cells (**fig 4, B**). We generated five virtual capture plots, examining all pairwise chromatin interactions throughout the tiled region in which individual, informative “viewpoints” participate: three CTCF sites (two flanking the α-globin sub-TAD, and one situated between the R1 and R2 enhancers), as well as the R2 enhancer, and the α-globin promoters. Since the two α-globin promoters are identical in sequence save for a single SNP; we considered both as viewpoints simultaneously. As well as generating virtual capture plots from WT and R2-only erythroid cells, we re-analysed a previously published WT ES cell tiled-C dataset (Oudelaar et al., 2020), to serve as a non-erythroid control. Chromatin interactions between each CTCF site and the surrounding DNA were unperturbed in R2-only cells, demonstrating that the 65kb α-globin sub-TAD still forms in the absence of the R1, R3, Rm and R4 elements (**fig 4, C**). However, interrogation of R2’s chromatin interaction profile revealed a striking reduction in interaction frequency between R2 and the α-globin promoters, which was corroborated by reciprocal virtual capture from the promoters themselves (**fig 4, C**). Although the frequency of these interactions was reduced in R2-only erythroid cells from fetal livers, it was still significantly higher than that in the WT ES cell baseline.

### The α-SE constituents are not equivalent and perform two distinct functions

In the R2-only mouse model the R2 enhancer retains many characteristics of an active enhancer; however, its ability to recruit coactivators, interact with its target gene promoters, produce bi-directional eRNA transcripts, and up-regulate α-globin expression are all severely attenuated. We next set out to rebuild the α-SE in various configurations to determine the role of each of the other SE elements. Because rebuilding the SE entailed the generation of numerous genetic models – too many to reasonably study in mice – we moved back to the orthogonal *in vitro* embryoid body (EB)-based erythroid differentiation system (Francis et al., 2022). This allowed rapid engineering of hemizygous WT and R2-only ES cells, followed by production of genetically engineered mouse erythroid cells, in which we could study the activity of the α-SE.

Our initial conclusion that the α-SE combines additively was based on deleting individual elements from an otherwise intact SE (Hay et al., 2016). Therefore, to revalidate these findings we reconstituted those same deletions in hemizygous mouse ES cells. Analysis of chromatin accessibility and α-globin gene expression in EB-derived erythroid cells were entirely consistent with our previous findings (**fig 5, A, D**). Individual deletion of R1 (ΔR1) or R2 (ΔR2) significantly reduced α-globin expression, and deleting both (ΔR1R2), leaving only the R3, Rm and R4 elements, reduced expression to ∼2% of WT. Meanwhile, deleting R3 (ΔR3) or Rm (ΔRm) alone had no discernible effect on gene expression, and deleting R4 (ΔR4) led to a small (∼15%), but statistically significant reduction in α-globin expression.

**Figure 5.**
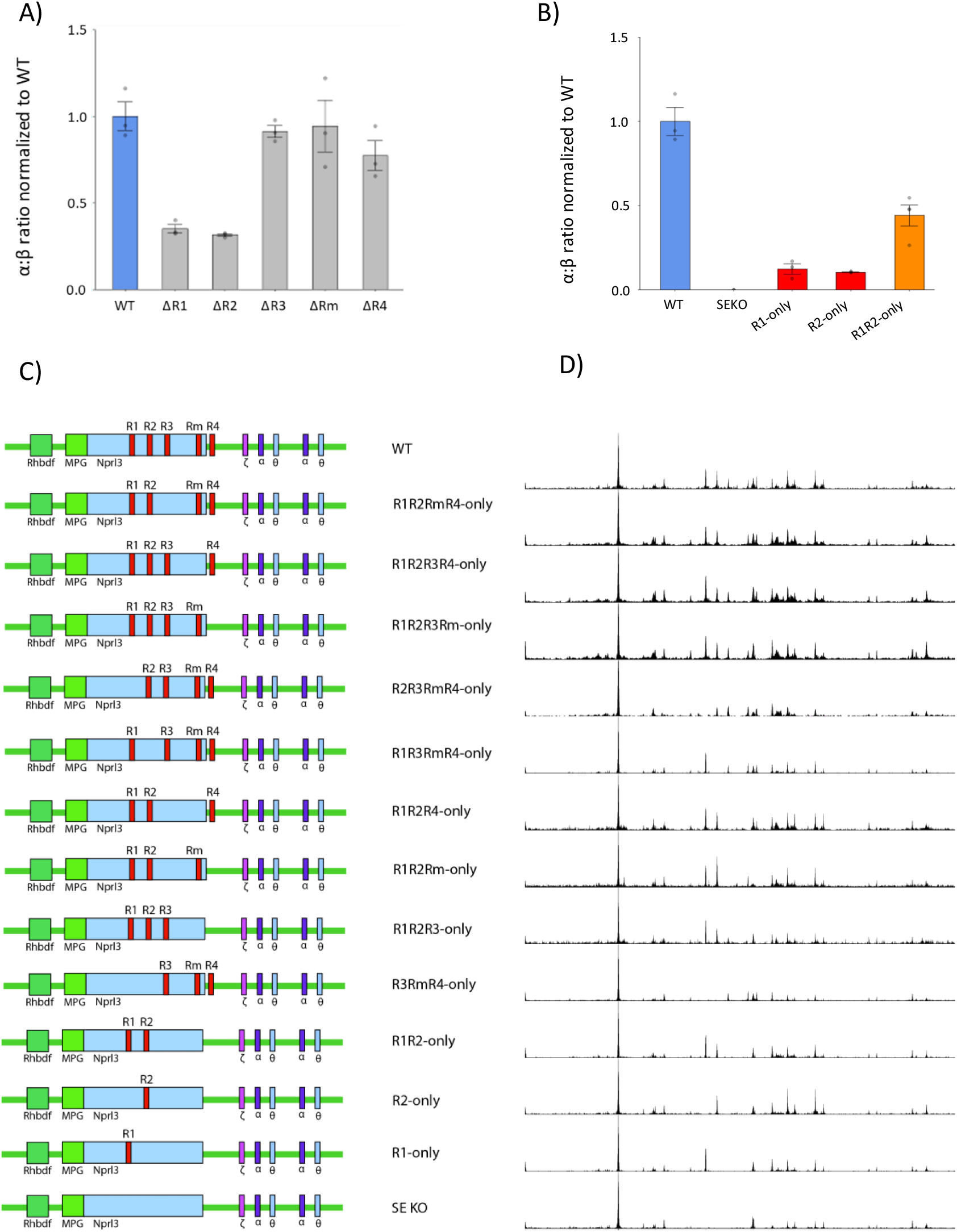
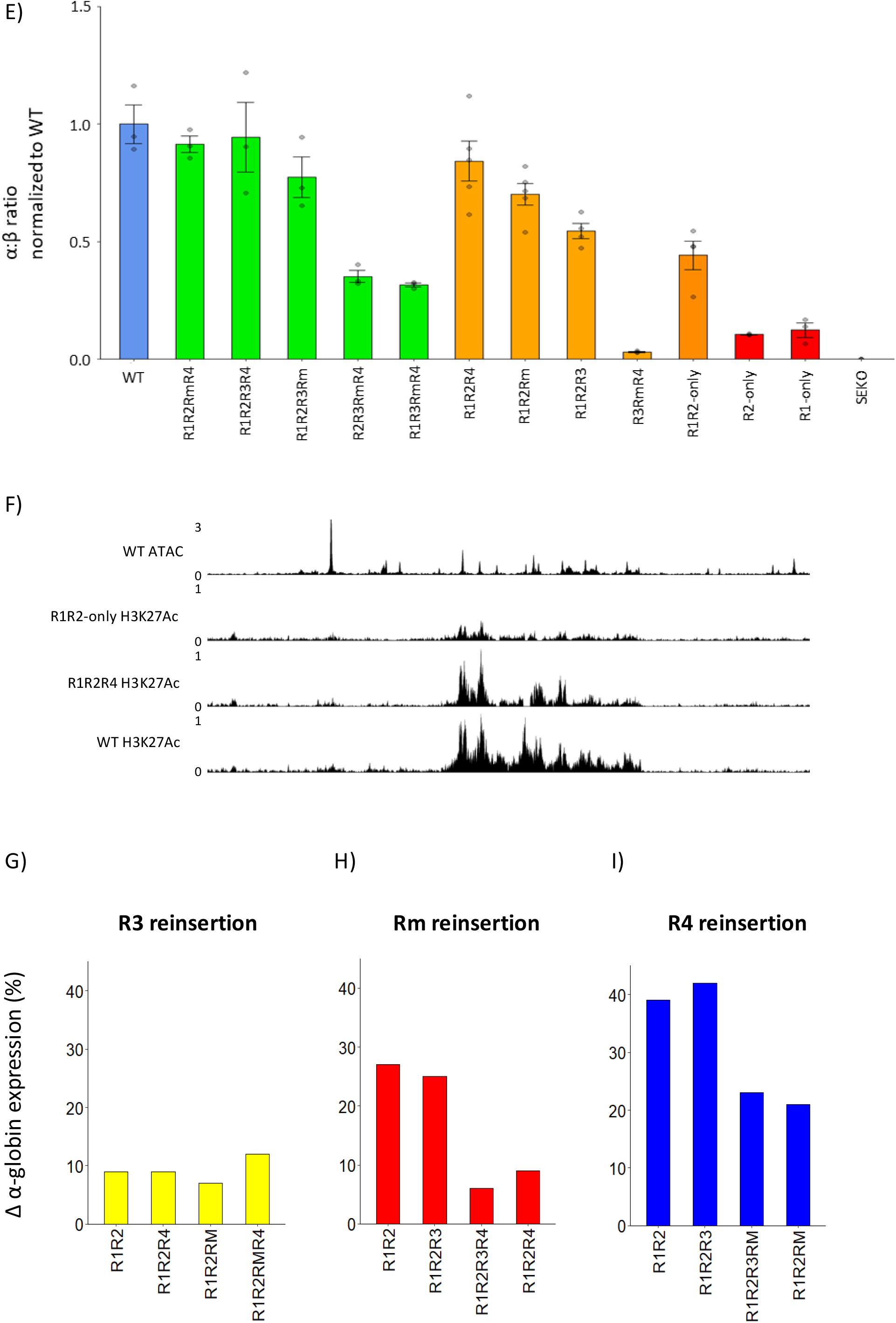
The R1 and R2 enhancers rely on R3, Rm and R4 in order to exert their full potential. A) EB-derived single deletion models recapitulate the results obtained in primary mouse erythroid cells. α-globin gene expression in WT, ΔR1, ΔR2, ΔR3, ΔRm and ΔR4 EB-derived erythroid cells (n≥3) assayed by RT-qPCR. Expression normalized to β-globin and displayed as a proportion of WT expression. Dots = biological replicates; error bars = SE. B) α-globin gene expression in WT, superenhancer knockout (SEKO), R1-only, R2-only and R1R2-only EB-derived erythroid cells (n≥3) assayed by RT-qPCR. Expression normalized to β-globin and displayed as a proportion of WT expression. Dots = biological replicates; error bars = SE. C) Graphical representation of the enhancer titration: fourteen genetic models rebuilding the α-SE in hemizygous mESCs. All models screened by PCR, Sanger sequencing and ATAC-seq. D) ATAC-seq in EB-derived erythroid cells in the corresponding enhancer titration models (n≥3). Tracks = merged biological replicates. E) α-globin gene expression in enhancer titration EB-derived erythroid cells (n≥3) assayed by RT-qPCR. Expression normalized to β-globin and displayed as a proportion of WT expression. Dots = biological replicates; error bars = SE. F) ATAC-seq in WT EB-derived erythroid cells (n=3, merged) (top). H3K27Ac ChIPmentation in R1R2-only (n=1), R1R2R4 (n=1) and WT (n=1) EB-derived erythroid cells. G) Percentage increase in α-globin expression following R3-reinsertion in each corresponding genetic background, as assayed by RT-qPCR in EB-derived erythroid cells. H) Percentage increase in α-globin expression following Rm-reinsertion in each corresponding genetic background, as assayed by RT-qPCR in EB-derived erythroid cells. I) Percentage increase in α-globin expression following R4-reinsertion in each corresponding genetic background, as assayed by RT-qPCR in EB-derived erythroid cells.

Next, we investigated whether reinserting the α-SE’s second major activator, R1, into the enhancerless Δα-SE locus would be capable of driving high levels of α-globin transcription. Similar to R2-only, EB-derived R1-only erythroid cells only expressed 10% α-globin compared to WT (**fig 5, B**). To our surprise, even a model harboring both major activators in their native positions (R1R2-only) was incapable of restoring high levels of α-globin transcription (**fig 5, B**).

The R3, Rm and R4 elements display little or no inherent conventional enhancer activity, but still they appear necessary for full α-SE activity. To investigate how R3, Rm and R4 complement the activity of R1 and R2, we generated an “enhancer titration series”, sequentially rebuilding the native α-SE from the deficient R1R2-only model to WT, generating all R3/Rm/R4 permutations (**fig 5, C**).

We generated at least three separately targeted clones for each model, and verified the integrity of each, using PCR and Sanger sequencing. To confirm that the newly designed models do not inadvertently create a sequence with potential function, we used the JASPAR and SASQUATCH *in silico* tools to screen for predicted changes in TF motifs and DNA accessibility at the deletion and insertion sites. Using ATAC-seq, we show that in each model the chromatin associated with the appropriate elements becomes accessible in erythroid cells, and there were no unexpected changes in accessibility throughout the remainder of the locus (**fig 5, D**).

To probe each model’s ability to enhance α-globin expression we performed RT-qPCR (**fig 5, E**). Reinserting the R3 element into the R1R2-only background only rescued gene expression by ∼10%, whereas reinsertion of Rm, or R4 had a much larger effect, rescuing expression by 25% and 40%, respectively. R4 reinsertion was accompanied by a large increase in H3K27 acetylation over the R1 and R2 elements (**fig 5, F**), suggesting that R4’s main role is to facilitate the full activity of the two major activators.

R3 reinsertion up-regulated α-globin transcription to approximately the same degree regardless of Rm and R4 coincidence (**fig 5, G**). Meanwhile, reintroducing Rm into a cluster containing R4 (e.g. inserting Rm into R1R2R4-only cells) only raised expression by 5-10% (**fig 5, H**). Likewise, the positive effect of reinserting R4 into a locus already containing Rm (e.g. reinserting R4 into R1R2Rm-only cells) was less than reinserting R4 into a locus containing only R1, R2 and/or R3 **(5, I**). Therefore, in their native context Rm and R4, but not R3, appear to be at least partially redundant in their ability to facilitate the function of the strong activators R1 and R2, with R4 having a stronger effect than Rm.

### R4’s rescue potential is derived from its position, and not its sequence

To investigate the cause of R4’s superior rescue potential, we re-analyzed an existing DNase-seq dataset, and conducted FIMO (MEME-suite) motif analysis on the five α-SE elements. Unsurprisingly, R1 and R2 contained the highest density of TF motifs, and the most complex DNase foot-printing signals. However, motif analysis demonstrated that R3 contains more erythroid TF motifs (absolute number and motif diversity) than Rm and R4 combined, which was supported by R3’s richer DNase foot-printing signal compared to Rm and R4 (**fig 6, A**). Inspection of Gata1, Nf-e2 and Tal1 ChIP-seq and ChIPmentation tracks from a number of WT erythroid tissues further supported the results of our motif analysis (data not shown). It is possible that R4 recruits other unknown factors, but our data suggest that the relative rescue capacities of R3, Rm and R4 are not primarily encoded in their relative capacities to recruit transcription factors.

**Figure 6.**
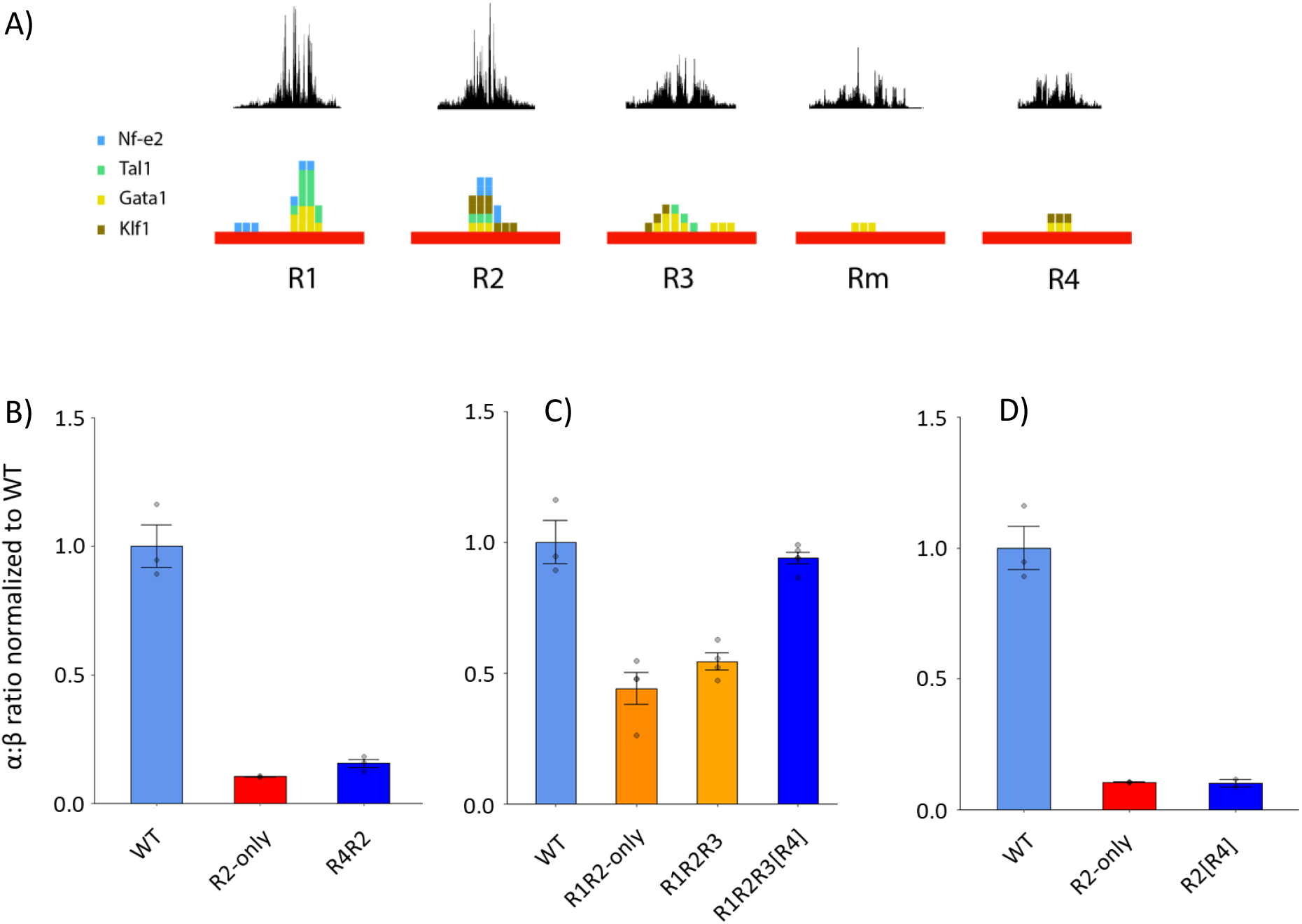
The relative activity of R3, Rm and R4 is primarily encoded in their positions rather than their sequences. A) Dnase foot-printing over each of the α-SE constituents (top). FIMO motif analysis conducted on each α-SE constituent, searching for occurrences of Gata1, Nf-e2, Tal1 and Klf1 motifs (bottom). B) α-globin gene expression in WT, R2-only and R4R2 EB-derived erythroid cells (n≥3) assayed by RT-qPCR. Expression normalized to β-globin and displayed as a proportion of WT expression. Dots = biological replicates; error bars = SE. C) α-globin gene expression in WT, R1R2-only, R1R2R3 and R1R2R3[R4] EB-derived erythroid cells (n≥3) assayed by RT-qPCR. Expression normalized to β-globin and displayed as a proportion of WT expression. Dots = biological replicates; error bars = SE. D) α-globin gene expression in WT (n=3), R2-only (n=3) and R2[R4] (n=2) EB-derived erythroid cells assayed by RT-qPCR. Expression normalized to β-globin and displayed as a proportion of WT expression. Dots = biological replicates; error bars = SE.

Rescue potential of R3, Rm and R4 inversely correlates with distance to the α-globin promoters. We therefore asked whether each element’s ability to bolster transcription depends more on its sequence or its proximity to the α-globin promoters. To test whether R4’s sequence is sufficient to rescue expression, we modified the R2-only model by reinserting R4; however, rather than placing R4 in its native position (close to the α-globin promoters), we reinserted it in the position of R1 (the element located furthest from the α-globin promoters) (R4R2). EB-derived R4R2 erythroid cells only expressed 12% α-globin, suggesting that R4’s rescue capacity is not exclusively based on its sequence (**fig 6, B**).

Next, to test the importance of element positioning, we modified the R1R2-only model, by inserting R3 in the position of R4 (R1R2R3[R4]). Moving R3 closer to the α-globin promoters in this manner had a dramatic effect, increasing gene expression by 50%, compared to the 10% rescue driven by R3 in its native position. Together, this strongly indicates that R4’s position, rather than its sequence, underpins its potency in rescuing gene expression (**fig 6, C**).

R2-only FL cells exhibited reduced interaction frequency between R2 and the α-globin promoters, and it seems that R4’s position, close to the α-globin promoters, is important for facilitating full R1 and R2 enhancer activity. We therefore speculated that R4 might play a role in increasing interaction frequency or stability between the α-SE and promoters. To test whether the R2-only transcriptional deficit could be rescued by simply reducing the linear distance between R2 and its cognate promoters, we modified the Δα-SE model, inserting R2 at the position of R4 (R2[R4]).

To our surprise, moving R2 closer to the α-globin promoters had no positive effect on gene expression (**fig 6, D**). This demonstrates that the physical linear proximity of R2 to the α-globin promoter was insufficient to restore R2’s full activity. Still, R4 could stabilize interactions, or aid the formation of particular 3D chromatin structures.

## Discussion

Since the seminal description of an enhancer element in 1981 (Banerji et al., 1981) and two years later the first report of what was effectively an enhancer cluster (Mercola et al., 1983), there has been an immense amount of research into what enhancers are, how they work and how they influence development and disease. Despite the fact that enhancer clusters have been studied for nearly forty years, we are yet to understand many of the most basic principles governing their activity, from the manner(s) by which cluster constituents cooperate with one another, to the biochemical processes compelling target gene up-regulation.

Over the years many groups have reported different “flavours” of biologically significant enhancer clusters, among them: locus control regions (Grosveld et al., 1987), shadow enhancers (Hong et al., 2008), regulatory archipelagos (Montavon et al., 2011), Greek islands (Markenscoff-Papadimitriou et al., 2014), stretch enhancers (Parker et al., 2013) and super-enhancers (Whyte et al., 2013). Many enhancer clusters satisfy the criteria of multiple classes. To simplify our analyses, we focused on studying the functional characteristics of SEs, selecting this particular class due to the clear bioinformatic definition of SEs (using the ROSE algorithm) and the fact that the field has widely adopted the “super-enhancer” nomenclature. SEs are defined by high levels of enhancer-associated H3K27Ac, high levels of TF and Mediator occupancy and the limited genomic distances between their constituents (Whyte et al., 2013). Numerous publications have demonstrated that SEs activate high levels of gene expression, with a tendency to regulate lineage-specific genes (Grosveld et al., 2021; Hnisz et al., 2015; Wang et al., 2019; Whyte et al., 2013). Despite this, it remains unclear whether there is a clear *functional* distinction separating SEs from clusters of regular enhancer elements. Perhaps *the* key question is whether SEs are clusters of independent elements combining in an additive fashion, or cohesive units exhibiting activities greater than the sum of their parts (Blobel et al., 2021; Grosveld et al., 2021). Recently, a number of groups have reported examples of SEs which appear to combine in non-additive fashions, presenting evidence of super-additive (Thomas et al., 2021), redundant (Hörnblad et al., 2021), synergistic (Shin et al., 2016), and hierarchical (Huang et al., 2018) cooperation between constituents. The majority of such studies have been confounded by either incomplete dissection of their model clusters, or disruption of clusters which regulate pleiotropic transcription factors or co-factors that influence cell fate, rendering it impossible to control whether WT and manipulated models are equivalent in their developmental stage and cell type.

Here, we have comprehensively dissected the tractable α-globin SE *in situ* to investigate how its five constituent elements cooperate. The α-SE is an ideal genetic model for this study; the SE and the TAD in which it is contained have been extensively characterized (Oudelaar et al., 2021), and the SE is activated exclusively during terminal erythroid differentiation, meaning its manipulation has no effect in non-erythroid cells. Previous dissection of the α-SE suggested that its five constituents combine additively as independent elements (Hay et al., 2016), a conclusion drawn through generating a series of mouse models harboring single and selective pairwise element deletions from an otherwise intact SE. Our present work further evaluates this conclusion, and demonstrates unequivocally that R2 requires (a subset of) the other four α-SE constituents to achieve its full enhancer potential. Despite maintaining the bioinformatic signature of an active enhancer, R2 by itself is not sufficient to up-regulate high levels of α-globin expression, exhibiting very low levels of co-activator recruitment, reduced interactions with its target genes’ promoters, and lower levels of eRNA transcription.

Rebuilding the α-SE from the Δα-SE model demonstrated that our previous single deletion models were simply inadequate to fully resolve the cooperation between the five α-SE constituents. This serves as a cautionary tale and clearly shows that extensive genetic dissection is essential to fully understand how an enhancer cluster operates. Combinatorial reconstruction of the α-SE exposed a complex network of functional interactions between its constituents: R1 and R2 cooperate synergistically, each up-regulating gene expression 100-fold, versus 450-fold when combined, whereas the R3, Rm and R4 elements display no intrinsic enhancer activity; instead, they facilitate the activities of R1 and R2.

Of note, the three non-enhancer “facilitator” elements display a hierarchy wherein R4 is the most potent facilitator and R3 the least. Whereas R3 facilitates the activities of R1 and R2 to a similar degree regardless of Rm/R4 coincidence, Rm and R4 function in a context-dependent manner, each partially redundant to the other. Interestingly, Sahu and colleagues recently used a STARR-seq method to show that four out of the five MYC SE constituents have no detectable enhancer activity in HepG2 cells (Sahu et al., 2022). It is unclear whether these four non-functional elements are required for full SE activity, but it raises the possibility that facilitators could be a common feature of SEs.

The mechanism(s) via which facilitators augment SE activity remain elusive, although it seems that their hierarchy is encoded in their positions rather than their sequences. Moving R2 closer to the α-globin promoters had no effect on gene expression, suggesting that facilitators do not solely act to increase enhancer-promoter interaction frequency; nevertheless, this does not preclude them playing a role in forming or stabilizing *specific* 3-dimensional structures. A recent interesting study in *Drosophila* has identified what may be similar elements which are not enhancers but are thought to facilitate interactions between regulatory elements by tethering them together (Levo et al., 2022).

A number of studies have suggested that enhancer clusters, including SEs, may act cooperatively to form dense foci containing high concentrations of transcriptional apparatus, including tissue-specific TFs, coactivators such as the mediator complex, and PolII (Grosveld et al., 2021). The biochemical processes leading to formation of such subnuclear structures is debated, but two prominent theories include liquid-liquid phase separation (Boija et al., 2018; Gurumurthy et al., 2019; Hnisz et al., 2017; Sabari et al., 2018) and some form of TF trapping (Sigova et al., 2015; Thomas et al., 2021). Both of these proposals require recruitment of a critical mass of TFs within a given 3 -dimensional space. It is feasible that R4 could rescue gene expression by increasing the density of TFBSs at a particularly influential position along the chromatin fibre; subsequent recruitment of coactivators such as the mediator complex could then be instructive for establishing a regulatory hub. Though speculative, this explanation is consistent both with R3’s ability to rescue transcription when transplanted to the position of R4 in the R1R2R3[R4] model, and R2’s continued insufficiency in the R2[R4] model.

In summary, our findings demonstrate that SEs can constitute complex cohesive networks of regulatory elements, displaying simultaneous additive, redundant, and synergistic cooperation. We present evidence that SEs can act as cohorts of functionally distinct elements, including activators, responsible for activating a target gene’s expression, and facilitators which facilitate efficient activator function. Without facilitators we see severely attenuated coactivator recruitment, enhancer-promoter interaction frequency, and eRNA transcription. Most importantly, we rigorously show that SEs do manifest emergent properties distinguishing them from clusters of regular enhancer elements.

## Materials and methods

### Synthetic BAC generation

Two RMGR-ready versions of the α-globin locus, the first encoding the five enhancer elements and the second deleting all but the R2 element, were constructed. A previously constructed BAC spanning the α-globin locus plus RMGR parts (Wallace et al., 2007) (RP23-46918; BACPAC Resources Centre, Children’s Hospital Oakland Research Institute; (Osoegawa et al., 2000)) was used as template to generate PCR amplicons with 50-200 base pairs of overlapping sequence for yeast homologous recombination. Gblocks (IDT) or fusion PCR products were used to provide homology with non-overlapping adjacent segments (e.g. enhancer deletions, vector-adjacent amplicons). A variant of the eSwAP-In method (Mitchell et al., 2021) was used to produce the two constructs, which were sequence verified using Illumina short read sequencing (**supplementary fig 1, A**).

### BAC transfection

RMGR-competent mouse ES cells were co-transfected by lipofection with Purified BAC DNA and a Pcaggs-Cre-IRES*puro* plasmid (A. J. H. Smith et al., 2002). Cells were selected for *Hprt* complementation, and the *Hprt* gene later removed by transfection with a transient flippase-expressing plasmid (Schaft et al., 2001). Cells were screened by selection with 6-thioguanine and PCR.

The structural integrity of the genome-integrated, BAC-derived R2-only locus was screened using 10X linked-read sequencing (**supplementary fig. 1, B**). Subsequent BAC-integrated loci were screened by PCR, Sanger sequencing, and ATAC-seq.

### CRISPR-Cas9 editing

Guide RNA (sequences in **supplementary table 2**) design and cloning were performed by the Weatherall Institute of Molecular Medicine genome engineering facility. Guide RNAs were designed using the CRISPOR and BreakingCas online gRNA design tools. Candidates with the fewest predicted off-targets were selected and further screened for their effectiveness, using an *in vitro* surveyor assay (according to the manufacturer, IDT).

For enhancer deletion, gRNAs were designed flanking the targeted enhancer, and cloned into pSpCas9 (BB)-2A-GFP (pX458) vector, a gift from Feng Zhang (Addgene plasmid: #48138), or pX458-ruby (Kredel et al., 2009)). Hemizygous WT mESCs were co-transfected, by lipofection, with the appropriate 5’ targeting vector (expressing GFP) and 3’-targeting vector (expressing mRuby), and 24-36 hours later, GFP-mRuby co-fluorescent cells were FACS sorted into individual wells of a 96 well plate. Individual clones were grown in each well for 8-10 days without disruption. When colonies were visible in each well, cells were split into two plates: one for screening, and the other for analysis/freezing. Clones were screened for successful enhancer deletion by boundary PCR; this entailed PCR amplification using primers flanking each deleted element (sequences in **supplementary table 3**), such that a successfully deleted allele would produce a smaller product than a WT allele. Clones were then screened by Sanger sequencing and ATAC-seq.

To produce R2[R4] and R1R2R3[R4] mutants, existing hemizygous Δα-SE and R1R2-only mutants were re-targeted. Each new model was generated using a single round of targeting – either through insertion of the R2 element at the position of R4 in Δα-SE cells, or insertion of the R3 element in the position of R4 in R1R2-only cells. Homology directed repair (HDR) donors were designed encoding the R2 element flanked by 500bp homology arms, homologous to the native position of the R4 element. A Sal1 restriction enzyme recognition site was inserted at the 5’ of the R2 enhancer in the HDR donor, and an Mlu1 site at the 3’ of the element. This enabled efficient restriction-ligation exchange of the R2 element within the HDR donor with the R3 element. The HDR donor construct was ordered as a GeneART Gene synthesis custom design. The HDR donor was also designed to inactivate the protospacer adjacent motif. R2[R4] and R1R2R3[R4] donors were screened with sasquatch and jaspar to ensure no novel accessibility sites or motifs were predicted (Fornes et al., 2020; Schwessinger et al., 2017), prior to synthesis and transfection.

Δα-SE and R1R2-only cells were co-transfected, by lipofection, with pX458 vectors (expressing gRNA targeting the position of R4) and the appropriate HDR donor. 24-36 hours post-transfection, GFP positive cells were FACS sorted into single wells in a 96-well format, and screened as described.

### Mouse model generation

All mouse work was performed in accordance with UK Home office regulations, under the appropriate animal licenses. Mouse model generation and animal husbandry was conducted by the Mouse Transgenics Core Facility at the Weatherall Institute of Molecular Medicine. R2-only BAC-integrated mESCs were karyotyped, microinjected into C57BL/6 blastocysts and implanted into pseudo-pregnant females. Chimeric males were back-crossed with WT females, and pups were screened by PCR for germline transmission.

Timed-heterozygote crosses: R2-only homozygotes were not viable, and therefore all analyses were restricted to embryonic timepoints. Pregnant mice were sacrificed at embryonic days E8.5, E9.5, E10.5, E12.5, E14.5, or E17.5 post-plug. Embryos were dissected from the pregnant females, ordered based on their predicted genotypes, and photographed. Erythropoietic cells/compartments were then isolated for analysis. Whole E8.5-10.5 embryos were mechanically disaggregated in heparinised PBS, and erythroid-containing supernatant aspirated into fresh tubes for processing; remaining material was stored for genotyping by PCR. Foetal livers (the definitive erythroid compartment) were isolated from E12.5-E17.5 embryos. Foetal livers were mechanically disaggregated to a single cell suspension in FACS buffer, and filtered through pre-separation filters; brain tissue was stored for genotyping by PCR and gene expression analysis (RT-qPCR). Erythroid cells were processed for analysis by RT-qPCR/RNA-seq, ATAC-seq, ChIP/ChIPmentation and 3C-based methods on the day of harvest (see below). FACS analysis, staining for the CD71 and Ter119 cell surface markers in E12.5 foetal liver cells, revealed that WT and R2-only foetal livers are composed of ∼95% CD71+/Ter119+ erythroid cells, indicating that no further selection (beyond mechanical disaggregation, and filtration through pre-separation filters (miltenyibiotec)) was required.

### Cell culture and in vitro erythroid differentiation system

E14-TG2a.IV (E14) mESCs, or genetic models derived from these cells, were cultured in gelatinised plates using standard methods (Jackson et al., 2010; A. G. Smith, 1991): cells were maintained in ES-complete medium, a GMEM-based medium supplemented with Leukemia inhibitory factor (LIF).

An in vitro EB-based differentiation system was used to generate erythroid cells (Francis et al., 2022). Briefly, 24 hours pre-differentiation, mouse ES cells were passaged into adaptation medium. mESCs transferred into 10cm petri dishes containing differentiation media (lacking LIF, and supplemented with transferrin) and cultured for seven days. After seven days of differentiation, CD71+ erythroid cells were selected and isolated. EBs were disaggregated into single cell suspension, through incubation in 0.25% trypsin for ∼3 minutes, and then quenched with FCS-containing media. Cells were incubated with anti-CD71 FITC-conjugated antibodies (Invitrogen, eBioscience 11-0711-85), followed by anti-FITC separation microbeads. CD71+ cells were then isolated by magnetic column separation (LS Column, Miltenyi), according to the manufacturer’s protocol.

### Gene expression analysis

On the day of cell harvest, aliquots of 5×105 cells (primary mouse cells, or CD71+ mouse ES cell-derived models) were lysed in trizol reagent, before being snap frozen and stored at -80°C. RNA was extracted using the Direct-zol MicroPrep kit (Zymo Research), according to the manufacturer’s protocol (however, the 15 minute DNase treatment step was lengthened to 45 minutes). RNA quality was assessed by tape station, using RNA screentape (Agilent). Only samples with an RNA integrity score of at least 8 were taken forwards for subsequent analysis. The extracted RNA was reverse transcribed using Superscript III First-Strand Synthesis SuperMix (Life Technologies).

qPCR with taqman probes (**supplementary table 4**) was utilised to analyse gene expression in each model. Results were normalised to RPS18 or the relevant β-globin genes. Analysis steps including all statistical tests (ANOVA) and graphical plotting were conducted in RStudio. The R package ggplot2 was used to generate and render each plot.

### NGS assays

#### ATAC-seq

Assay for transposase-accessible chromatin (ATAC)-seq was performed on ∼7×104 cells, using the Illumina Tagment DNA enzyme and buffer kit (illumina), as previously described (Buenrostro et al., 2015; Hay et al., 2016). Briefly, cells were lysed in a gentle NP-40 containing lysis buffer, and resuspended in Tn5 buffer with illumina adaptor-loaded Tn5 enzyme. Cells were incubated for 30 minutes at 37°C, and then tagmented DNA was purified using XP Ampure beads (mybeckman), before indexing with Nextera indexing primers (illumina). Indexed ATAC samples were assessed by tape station, using a D1000 HS screen tape (Agilent).

### ChIPmentation

ChIPmentation experiments were performed as previously described (Schmidl et al., 2015), with few modifications. Briefly, on the day of cell harvest, aliquots of 1×105-1×106 cells (primary mouse cells, or CD71+ mouse ES cell-derived models) were either single-fixed with 1% formaldehyde for 10 minutes, followed by quenching with 125mM glycine, or double-fixed with 2mM disuccinimidyl glutarate (DSG) for 50 minutes, followed by 1% formaldehyde for 10 minutes, before quenching with 125mM Glycine. Single-fixed samples were ultimately used for ChIPmentation experiments assaying histone modifications; double-fixed samples were used for experiments assaying transcription factor occupancy.

Cells were spun down and washed with PBS, before being snap frozen. Fixed aliquots were stored at -80°C. Cell pellets were lysed in 0.5% SDS lysis buffer and sonicated, using a covaris ME220 sonicator, to fragment DNA to an average fragment length of ∼2-300bp. Sonicated chromatin was analysed by tape station, using a D1000 or D1000 HS screen tape (Agilent). SDS in the lysis buffer was neutralised with 1% triton-X, and the sonicate was incubated overnight with a mix of protein A and G dynabeads (Thermofisher) and the appropriate antibody (**supplementary table 5**). The following morning, chromatin-bound beads were washed three times using a low salt, a high salt and a LiCl-containing wash buffer, followed by tagmentation of the immunoprecipitated chromatin with sequencing adaptor-loaded tn5. Samples were indexed, using Nextera indices (Illumina).

### Tiled-C

Tiled-C was conducted as previously described (Oudelaar et al., 2020). Briefly, on the day of harvest, aliquots of 5×10^5^ cells (primary mouse cells, or CD71+ mouse ES cell-derived models) were fixed with 2% formaldehyde for 10 minutes, before quenching with 125mM Glycine.

Cells were spun down, washed with PBS, and the pellet suspended in a mild NP-40-containing lysis buffer. Samples were then snap frozen and stored at -80°C. Cells in lysis buffer were thawed and spun down, before resuspension in restriction enzyme buffer mix. An appropriate volume of DpnII was added, and samples were incubated overnight at 37°C. Fresh aliquots of DpnII were added the following morning and afternoon. The DpnII was heat inactivated, and proximal DpnII-digested “sticky ends” were ligated using T4 ligase. Digested-re-ligated DNA was extracted using XP Ampure beads (mybeckman) and sonicated using a covaris ME220 sonicator. Sonicated chromatin was analysed by tape station, using a D1000 or D1000 HS screen tape (Agilent). The resultant fragments were indexed using the NEBNext Ultra II library preparation kit (New England BioLabs). Fragments corresponding to the region of interest (chr11:29902951-33226736) were enriched using oligo capture with biotinylated oligos (Twist) complementary to every DpnII fragment within the tiled region, before streptavidin pulldown using Streptavidin dynabeads (Thermofisher).

### RNA-seq

On the day of harvest, aliquots of 5×10^5^ cells (primary mouse cells, or CD71+ mouse ES cell-derived models) were lysed in trizol reagent, before being snap frozen and stored at -80°C. RNA was extracted using the Direct-zol MicroPrep kit (Zymo Research), according to the manufacturer’s protocol (however, the 15 minute DNase treatment step was lengthened to 45 minutes). RNA quality was assessed by tape station, using RNA screentape (Agilent). Only samples with an RNA integrity score of at least 8 were used.

Poly-A positive and negative RNA-seq was performed on 5×10^5^ cells, using the NEBNext Ultra II Directional RNA Library Prep Kit for Illumina (New England BioLabs). Ribosomal RNA was depleted using the NEBNext rRNA Depletion Kit, and then the poly-A positive and negative fractions were separated using the NEBNext Poly(A) mRNA Magnetic Isolation Module.

### Sequencing and bioinformatic analysis

All NGS sequencing was performed using TG NSQ 500/550 Hi Output v2.5 (75 CYS) kits (illumina); these kits are paired-end sequencing kits which produce two 40 base pair reads, corresponding to the 5’ and the 3’ of the fragment being sequenced. Generally, ∼25-40 million reads were desirable for each ATAC or ChIPmentation sample, ∼10-20 million for each RNA-seq sample, and ∼5-10 million for each Tiled-C sample, although actual sequencing depth was variable.

### ATAC and ChIPmentation

The quality of the FASTQ files from ATAC-seq and ChIPmentation were assessed using FASTQC, and the reads aligned to the mm9 mouse genome, using bowtie2. Non-aligning reads were trimmed using Cutadapt trimgalore and then realigned to the mm9 genome using bowtie2. All reads which still failed to align were extracted, and flashed using FLASh, before realignment to the mm9 genome using bowtie2. All of the files containing successfully aligning reads were concatenated, and aligned to the mm9 genome together using bowtie2. Resultant SAM files were filtered, sorted, and PCR duplicates removed, using SAMtools (samtools view, sort, and rmdup, respectively). The resultant BAM file was indexed using SAMtools index, and converted to a bigwig file using deeptools bamcoverage. Each bigwig was visualised using the University of California Santa Cruz (UCSC) genome browser, and traces corresponding to regions of interest were downloaded from here. Peaks were called in each sample using macs2 with default parameters, and differential accessibility/binding analysis was conducted using Bioconductor DESeq2 in RStudio. Motif analysis was performed using the MEME suite (meme-chip for de novo motif analysis and fimo for finding occurrences of known motifs), using HOCOMOCO mouse position weight matrices. Principal component analysis was performed on ATAC samples, using the DiffBind, rgl and magick packages in RStudio.

### Tiled-C

Tiled-C samples were analysed using the HiC-Pro pipeline, using the capture Hi-C workflow (aligning the data to the mm9 genome). To avoid interaction bias between regions within and outside of the tiled region, all data mapping to the tiled region was extracted and the remaining data discarded from subsequent analysis steps. Interaction matrices were ICE-normalised using HiC-Pro, and heatmaps generated for visualisation using ggplot2 in RStudio. Virtual capture plots were generated by extracting all entries within the tiled-C matrix in which a specific viewpoint of interest participates, and interaction scores normalised by dividing interaction scores by the total number of interactions within the tiled region. Virtual capture plots were produced for visualisation using ggplot2 in RStudio.

### RNA-seq

RNA-seq data was aligned to the mm9 genome, using star. The resultant SAM files were then filtered and sorted using SAMtools (samtools view and sort, respectively). The resultant BAM files were indexed using SAMtools index, and directional, rpkm normalised bigwigs generated using deeptools bamcoverage, with the filteredRNAstrand flag enabled. Each sample bigwig was visualised using the University of California Santa Cruz (UCSC) genome browser, and traces corresponding to regions of interest downloaded from here. Read coverage over each gene in the mm9 genome was calculated using Rsubread featurecounts, and differential expression analysis performed using edgeR in RStudio. Plots were generated using ggplot2 in Rstudio. Principal component analysis was performed on RNA-seq samples, using the DiffBind, rgl and magick packages in RStudio. To compare enhancer RNA transcription in WT and R2-only cells, levels of poly-A negative RNA over the R1, R2, R3, Rm and R4 enhancers were visually assessed on the UCSC genome browser; however, this was only possible on the + strand, as the Nprl3 gene, in which the R1, R2 and R3 enhancers are located, is transcribed on the – strand. To compare R2 enhancer RNA transcription quantitatively, a virtual qPCR was performed, by normalizing the number of reads mapping to the R2 enhancer in each sample to the number of reads mapping to the HS2 enhancer of the β-globin LCR or the RPS18 gene in the same sample. Levels of the normalised enhancer RNA transcription in WT and R2-only samples were then compared.

## Acknowledgements

The authors would like to express their gratitude to all colleagues who contributed to this work, in particular: Jackie Sloane-Stanley and the Weatherall Institute of Molecular medicine Transgenics core facility, Dr Philip Hublitz and the Weatherall Institute of Molecular medicine Genome engineering facility, and the Weatherall Institute of Molecular medicine flow cytometry facility.

## Funding statement

This study was not specifically funded. The main contributing authors are funded as detailed below:

Wellcome Trust: Joseph Blayney 219979/Z/19/Z

UKRI | Medical Research Council (MRC): MR/T014067/1

Wellcome Trust: Helena Francis 109097/Z/15/Z

Supported in part by grant RM1HG009491 to JDB.

Wellcome Trust: Rosa Stolper 215111/Z/18/Z

The funders had no role in study design, data collection and analysis, decision to publish, or preparation of the manuscript.

## References

Allahyar, A., Vermeulen, C., Bouwman, B. A. M., Krijger, P. H. L., Verstegen, M. J. A. M., Geeven, G., van Kranenburg, M., Pieterse, M., Straver, R., Haarhuis, J. H. I., Jalink, K., Teunissen, H., Renkens, I. J., Kloosterman, W. P., Rowland, B. D., de Wit, E., de Ridder, J., & de Laat, W. (2018). Enhancer hubs and loop collisions identified from single-allele topologies. Nature Genetics, 50(8), 1151–1160. https://doi.org/10.1038/S41588-018-0161-5

Arnold, P. R., Wells, A. D., & Li, X. C. (2020). Diversity and Emerging Roles of Enhancer RNA in Regulation of Gene Expression and Cell Fate. Frontiers in Cell and Developmental Biology, 7, 377. https://doi.org/10.3389/FCELL.2019.00377/BIBTEX

Banerji, J., Rusconi, S., & Schaffner, W. (1981). Expression of a beta-globin gene is enhanced by remote SV40 DNA sequences. Cell, 27(2 Pt 1), 299–308. https://doi.org/10.1016/0092-8674(81)90413-X

Beagrie, R. A., Scialdone, A., Schueler, M., Kraemer, D. C. A., Chotalia, M., Xie, S. Q., Barbieri, M., de Santiago, I., Lavitas, L. M., Branco, M. R., Fraser, J., Dostie, J., Game, L., Dillon, N., Edwards, P. A. W., Nicodemi, M., & Pombo, A. (2017). Complex multi-enhancer contacts captured by genome architecture mapping. Nature 2017 543:7646, 543(7646), 519–524. https://doi.org/10.1038/nature21411

Bender, M. A., Ragoczy, T., Lee, J., Byron, R., Telling, A., Dean, A., & Groudine, M. (2012). The hypersensitive sites of the murine β-globin locus control region act independently to affect nuclear localization and transcriptional elongation. Blood, 119(16). https://doi.org/10.1182/blood-2011-09-380485

Blobel, G. A., Higgs, D. R., Mitchell, J. A., Notani, D., & Young, R. A. (2021). Testing the super-enhancer concept. Nature Reviews. Genetics, 22(12), 749–755. https://doi.org/10.1038/S41576-021-00398-W

Boija, A., Klein, I. A., Sabari, B. R., Dall’Agnese, A., Coffey, E. L., Zamudio, A. v., Li, C. H., Shrinivas, K., Manteiga, J. C., Hannett, N. M., Abraham, B. J., Afeyan, L. K., Guo, Y. E., Rimel, J. K., Fant, C. B., Schuijers, J., Lee, T. I., Taatjes, D. J., & Young, R. A. (2018). Transcription Factors Activate Genes through the Phase-Separation Capacity of Their Activation Domains. Cell, 175(7), 1842–1855.e16. https://doi.org/10.1016/J.CELL.2018.10.042

Buenrostro, J. D., Wu, B., Chang, H. Y., & Greenleaf, W. J. (2015). ATAC-seq: A Method for Assaying Chromatin Accessibility Genome-Wide. Current Protocols in Molecular Biology / edited by Frederick M. Ausubel … [et Al.], 109, 21.29.1. https://doi.org/10.1002/0471142727.MB2129S109

Dębek, S., & Juszczyński, P. (2022). Super enhancers as master gene regulators in the pathogenesis of hematologic malignancies. Biochimica et Biophysica Acta (BBA) - Reviews on Cancer, 1877(2), 188697. https://doi.org/10.1016/J.BBCAN.2022.188697

Fornes, O., Castro-Mondragon, J. A., Khan, A., van der Lee, R., Zhang, X., Richmond, P. A., Modi, B. P., Correard, S., Gheorghe, M., Baranašić, D., Santana-Garcia, W., Tan, G., Chèneby, J., Ballester, B., Parcy, F., Sandelin, A., Lenhard, B., Wasserman, W. W., & Mathelier, A. (2020). JASPAR 2020: Update of the open-Access database of transcription factor binding profiles. Nucleic Acids Research, 48(D1). https://doi.org/10.1093/nar/gkz1001

Francis, H. S., Harold, C. L., Beagrie, R. A., King, A. J., Gosden, M. E., Blayney, J. W., Jeziorska, D. M., Babbs, C., Higgs, D. R., & Kassouf, M. T. (2022). Scalable in vitro production of defined mouse erythroblasts. PLOS ONE, 17(1), e0261950. https://doi.org/10.1371/JOURNAL.PONE.0261950

Grosveld, F., van Assendelft, G. B., Greaves, D. R., & Kollias, G. (1987). Position-independent, high-level expression of the human beta-globin gene in transgenic mice. Cell, 51(6), 975–985. https://doi.org/10.1016/0092-8674(87)90584-8

Grosveld, F., van Staalduinen, J., & Stadhouders, R. (2021). Transcriptional Regulation by (Super)Enhancers: From Discovery to Mechanisms. Annual Review of Genomics and Human Genetics, 22, 127–146. https://doi.org/10.1146/ANNUREV-GENOM-122220-093818

Gurumurthy, A., Shen, Y., Gunn, E. M., & Bungert, J. (2019). Phase Separation and Transcription Regulation: Are Super-Enhancers and Locus Control Regions Primary Sites of Transcription Complex Assembly? BioEssays, 41(1), 1800164. https://doi.org/10.1002/BIES.201800164

Hanssen, L. L. P., Kassouf, M. T., Oudelaar, A. M., Biggs, D., Preece, C., Downes, D. J., Gosden, M., Sharpe, J. A., Sloane-Stanley, J. A., Hughes, J. R., Davies, B., & Higgs, D. R. (2017). Tissue-specific CTCF-cohesin-mediated chromatin architecture delimits enhancer interactions and function in vivo. Nature Cell Biology, 19(8). https://doi.org/10.1038/ncb3573

Harteveld, C. L., & Higgs, D. R. (2010). Alpha-thalassaemia. Orphanet Journal of Rare Diseases, 5(1). https://doi.org/10.1186/1750-1172-5-13

Hay, D., Hughes, J. R., Babbs, C., Davies, J. O. J., Graham, B. J., Hanssen, L. L. P., Kassouf, M. T., Oudelaar, A. M., Sharpe, J. A., Suciu, M. C., Telenius, J., Williams, R., Rode, C., Li, P. S., Pennacchio, L. A., Sloane-Stanley, J. A., Ayyub, H., Butler, S., Sauka-Spengler, T., … Higgs, D. R. (2016). Genetic dissection of the α-globin super-enhancer in vivo. Nature Genetics, 48(8). https://doi.org/10.1038/ng.3605

Higgs, D. R., Engel, J. D., & Stamatoyannopoulos, G. (2012). Thalassaemia. Lancet (London, England), 379(9813), 373–383. https://doi.org/10.1016/S0140-6736(11)60283-3

Hnisz, D., Schuijers, J., Lin, C. Y., Weintraub, A. S., Abraham, B. J., Lee, T. I., Bradner, J. E., & Young, R. A. (2015). Convergence of Developmental and Oncogenic Signaling Pathways at Transcriptional Super-Enhancers. Molecular Cell, 58(2). https://doi.org/10.1016/j.molcel.2015.02.014

Hnisz, D., Shrinivas, K., Young, R. A., Chakraborty, A. K., & Sharp, P. A. (2017). A phase separation model predicts key features of transcriptional control. Cell, 169(1), 13. https://doi.org/10.1016/J.CELL.2017.02.007

Hong, J. W., Hendrix, D. A., & Levine, M. S. (2008). Shadow enhancers as a source of evolutionary novelty. Science (New York, N.Y.), 321(5894), 1314. https://doi.org/10.1126/SCIENCE.1160631

Hörnblad, A., Bastide, S., Langenfeld, K., Langa, F., & Spitz, F. (2021). Dissection of the Fgf8 regulatory landscape by in vivo CRISPR-editing reveals extensive intra- and inter-enhancer redundancy. Nature Communications, 12(1). https://doi.org/10.1038/s41467-020-20714-y

Hua, P., Badat, M., Hanssen, L. L. P., Hentges, L. D., Crump, N., Downes, D. J., Jeziorska, D. M., Oudelaar, A. M., Schwessinger, R., Taylor, S., Milne, T. A., Hughes, J. R., Higgs, D. R., & Davies, J. O. J. (2021). Defining genome architecture at base-pair resolution. Nature, 595(7865). https://doi.org/10.1038/s41586-021-03639-4

Huang, J., Li, K., Cai, W., Liu, X., Zhang, Y., Orkin, S. H., Xu, J., & Yuan, G. C. (2018). Dissecting super-enhancer hierarchy based on chromatin interactions. Nature Communications, 9(1). https://doi.org/10.1038/s41467-018-03279-9

Hughes, J. R., Cheng, J. F., Ventress, N., Prabhakar, S., Clark, K., Anguita, E., de Gobbi, M., de Jong, P., Rubin, E., & Higgs, D. R. (2005). Annotation of cis-regulatory elements by identification, subclassification, and functional assessment of multispecies conserved sequences. Proceedings of the National Academy of Sciences of the United States of America, 102(28), 9830–9835. https://doi.org/10.1073/PNAS.0503401102/SUPPL_FILE/03401FIG7.PDF

Hughes, J. R., Roberts, N., Mcgowan, S., Hay, D., Giannoulatou, E., Lynch, M., de Gobbi, M., Taylor, S., Gibbons, R., & Higgs, D. R. (2014). Analysis of hundreds of cis-regulatory landscapes at high resolution in a single, high-throughput experiment. Nature Genetics, 46(2). https://doi.org/10.1038/ng.2871

Ing-Simmons, E., Seitan, V. C., Faure, A. J., Flicek, P., Carroll, T., Dekker, J., Fisher, A. G., Lenhard, B., & Merkenschlager, M. (2015). Spatial enhancer clustering and regulation of enhancer-proximal genes by cohesin. Genome Research, 25(4), 504–513. https://doi.org/10.1101/GR.184986.114

Jackson, M., Taylor, A. H., Jones, E. A., & Forrester, L. M. (2010). The culture of mouse embryonic stem cells and formation of embryoid bodies. Methods in Molecular Biology (Clifton, N.J.), 633, 1–18. https://doi.org/10.1007/978-1-59745-019-5_1

King, A. J., Songdej, D., Downes, D. J., Beagrie, R. A., Liu, S., Buckley, M., Hua, P., Suciu, M. C., Marieke Oudelaar, A., Hanssen, L. L. P., Jeziorska, D., Roberts, N., Carpenter, S. J., Francis, H., Telenius, J., Olijnik, A. A., Sharpe, J. A., Sloane-Stanley, J., Eglinton, J., … Babbs, C. (2021). Reactivation of a developmentally silenced embryonic globin gene. Nature Communications 2021 12:1, 12(1), 1–15. https://doi.org/10.1038/s41467-021-24402-3

Kredel, S., Oswald, F., Nienhaus, K., Deuschle, K., Röcker, C., Wolff, M., Heilker, R., Nienhaus, G. U., & Wiedenmann, J. (2009). mRuby, a bright monomeric red fluorescent protein for labeling of subcellular structures. PLoS ONE, 4(2). https://doi.org/10.1371/journal.pone.0004391

Levo, M., Raimundo, J., Bing, X. Y., Sisco, Z., Batut, P. J., Ryabichko, S., Gregor, T., & Levine, M. S. (2022). Transcriptional coupling of distant regulatory genes in living embryos. Nature 2022 605:7911, 605(7911), 754–760. https://doi.org/10.1038/s41586-022-04680-7

Li, Y., Hu, M., & Shen, Y. (2018). Gene regulation in the 3D genome. Human Molecular Genetics, 27(R2), R228–R233. https://doi.org/10.1093/HMG/DDY164

Markenscoff-Papadimitriou, E., Allen, W. E., Colquitt, B. M., Goh, T., Murphy, K. K., Monahan, K., Mosley, C. P., Ahituv, N., & Lomvardas, S. (2014). Enhancer interaction networks as a means for singular olfactory receptor expression. Cell, 159(3), 543. https://doi.org/10.1016/J.CELL.2014.09.033

Mercola, M., Wang, X. F., Olsen, J., & Calame, K. (1983). Transcriptional enhancer elements in the mouse immunoglobulin heavy chain locus. Science, 221(4611), 663–665. https://doi.org/10.1126/SCIENCE.6306772

Mitchell, L. A., McCulloch, L. H., Pinglay, S., Berger, H., Bosco, N., Brosh, R., Bulajić, M., Huang, E., Hogan, M. S., Martin, J. A., Mazzoni, E. O., Davoli, T., Maurano, M. T., & Boeke, J. D. (2021). De novo assembly and delivery to mouse cells of a 101 kb functional human gene. Genetics, 218(1). https://doi.org/10.1093/GENETICS/IYAB038

Montavon, T., Soshnikova, N., Mascrez, B., Joye, E., Thevenet, L., Splinter, E., de Laat, W., Spitz, F., & Duboule, D. (2011). A regulatory archipelago controls Hox genes transcription in digits. Cell, 147(5), 1132–1145. https://doi.org/10.1016/J.CELL.2011.10.023

Moorthy, S. D., Davidson, S., Shchuka, V. M., Singh, G., Malek-Gilani, N., Langroudi, L., Martchenko, A., So, V., Macpherson, N. N., & Mitchell, J. A. (2017). Enhancers and super-enhancers have an equivalent regulatory role in embryonic stem cells through regulation of single or multiple genes. Genome Research, 27(2). https://doi.org/10.1101/gr.210930.116

Osoegawa, K., Tateno, M., Woon, P. Y., Frengen, E., Mammoser, A. G., Catanese, J. J., Hayashizaki, Y., & de Jong, P. J. (2000). Bacterial artificial chromosome libraries for mouse sequencing and functional analysis. Genome Research, 10(1). https://doi.org/10.1101/gr.10.1.116

Oudelaar, A. M., Beagrie, R. A., Gosden, M., de Ornellas, S., Georgiades, E., Kerry, J., Hidalgo, D., Carrelha, J., Shivalingam, A., El-Sagheer, A. H., Telenius, J. M., Brown, T., Buckle, V. J., Socolovsky, M., Higgs, D. R., & Hughes, J. R. (2020). Dynamics of the 4D genome during in vivo lineage specification and differentiation. Nature Communications, 11(1). https://doi.org/10.1038/s41467-020-16598-7

Oudelaar, A. M., Beagrie, R. A., Kassouf, M. T., & Higgs, D. R. (2021). The mouse alpha-globin cluster: a paradigm for studying genome regulation and organization. Current Opinion in Genetics & Development, 67, 18–24. https://doi.org/10.1016/J.GDE.2020.10.003

Oudelaar, A. M., Davies, J. O. J., Hanssen, L. L. P., Telenius, J. M., Schwessinger, R., Liu, Y., Brown, J. M., Downes, D. J., Chiariello, A. M., Bianco, S., Nicodemi, M., Buckle, V. J., Dekker, J., Higgs, D. R., & Hughes, J. R. (2018). Single-allele chromatin interactions identify regulatory hubs in dynamic compartmentalized domains. Nature Genetics, 50(12). https://doi.org/10.1038/s41588-018-0253-2

Oudelaar, A. M., Harrold, C. L., Hanssen, L. L. P., Telenius, J. M., Higgs, D. R., & Hughes, J. R. (2019). A revised model for promoter competition based on multi-way chromatin interactions at the α-globin locus. Nature Communications, 10(1). https://doi.org/10.1038/s41467-019-13404-x

Oudelaar, A. M., & Higgs, D. R. (2021). The relationship between genome structure and function. Nature Reviews Genetics, 22(3), 154–168. https://doi.org/10.1038/S41576-020-00303-X

Parker, S. C. J., Stitzel, M. L., Taylor, D. L., Orozco, J. M., Erdos, M. R., Akiyama, J. A., van Bueren, K. L., Chines, P. S., Narisu, N., Black, B. L., Axel, V., Pennacchio, L. A., & Collins, F. S. (2013). Chromatin stretch enhancer states drive cell-specific gene regulation and harbor human disease risk variants. Proceedings of the National Academy of Sciences of the United States of America, 110(44), 17921–17926. https://doi.org/10.1073/PNAS.1317023110/SUPPL_FILE/SD01.XLS

Pott, S., & Lieb, J. D. (2015). What are super-enhancers? Nature Genetics, 47(1), 8–12. https://doi.org/10.1038/NG.3167

Sabari, B. R., Dall’Agnese, A., Boija, A., Klein, I. A., Coffey, E. L., Shrinivas, K., Abraham, B. J., Hannett, N. M., Zamudio, A. v., Manteiga, J. C., Li, C. H., Guo, Y. E., Day, D. S., Schuijers, J., Vasile, E., Malik, S., Hnisz, D., Lee, T. I., Cisse, I. I., … Young, R. A. (2018). Coactivator condensation at super-enhancers links phase separation and gene control. Science (New York, N.Y.), 361(6400). https://doi.org/10.1126/SCIENCE.AAR3958

Sahu, B., Hartonen, T., Pihlajamaa, P., Wei, B., Dave, K., Zhu, F., Kaasinen, E., Lidschreiber, K., Lidschreiber, M., Daub, C. O., Cramer, P., Kivioja, T., & Taipale, J. (2022). Sequence determinants of human gene regulatory elements. Nature Genetics 2022 54:3, 54(3), 283–294. https://doi.org/10.1038/s41588-021-01009-4

Sartorelli, V., & Lauberth, S. M. (2020). Enhancer RNAs are an important regulatory layer of the epigenome. Nature Structural & Molecular Biology 2020 27:6, 27(6), 521–528. https://doi.org/10.1038/s41594-020-0446-0

Schaft, J., Ashery-Padan, R., van Hoeven, F. der, Gruss, P., & Francis Stewart, A. (2001). Efficient FLP recombination in mouse ES cells and oocytes. Genesis, 31(1). https://doi.org/10.1002/gene.1076

Schmidl, C., Rendeiro, A. F., Sheffield, N. C., & Bock, C. (2015). ChIPmentation: Fast, robust, low-input ChIP-seq for histones and transcription factors. Nature Methods, 12(10). https://doi.org/10.1038/nmeth.3542

Schwessinger, R., Suciu, M. C., McGowan, S. J., Telenius, J., Taylor, S., Higgs, D. R., & Hughes, J. R. (2017). Sasquatch: Predicting the impact of regulatory SNPs on transcription factor binding from cell-and tissue-specific DNase footprints. Genome Research, 27(10). https://doi.org/10.1101/gr.220202.117

Shin, H. Y., Willi, M., Yoo, K. H., Zeng, X., Wang, C., Metser, G., & Hennighausen, L. (2016). Hierarchy within the mammary STAT5-driven Wap super-enhancer. Nature Genetics, 48(8). https://doi.org/10.1038/ng.3606

Sigova, A. A., Abraham, B. J., Ji, X., Molinie, B., Hannett, N. M., Guo, Y. E., Jangi, M., Giallourakis, C. C., Sharp, P. A., & Young, R. A. (2015). Transcription factor trapping by RNA in gene regulatory elements. Science (New York, N.Y.), 350(6263), 978. https://doi.org/10.1126/SCIENCE.AAD3346

Smith, A. G. (1991). Culture and differentiation of embryonic stem cells. Journal of Tissue Culture Methods 1991 13:2, 13(2), 89–94. https://doi.org/10.1007/BF01666137

Smith, A. J. H., Xian, J., Richardson, M., Johnstone, K. A., & Rabbitts, P. H. (2002). Cre-loxP chromosome engineering of a targeted deletion in the mouse corresponding to the 3p21.3 region of homozygous loss in human tumours. Oncogene, 21(29). https://doi.org/10.1038/sj.onc.1205530

Tang, F., Yang, Z., Tan, Y., & Li, Y. (2020). Super-enhancer function and its application in cancer targeted therapy. Npj Precision Oncology 2020 4:1, 4(1), 1–7. https://doi.org/10.1038/s41698-020-0108-z

Thomas, H. F., Kotova, E., Jayaram, S., Pilz, A., Romeike, M., Lackner, A., Penz, T., Bock, C., Leeb, M., Halbritter, F., Wysocka, J., & Buecker, C. (2021). Temporal dissection of an enhancer cluster reveals distinct temporal and functional contributions of individual elements. Molecular Cell, 81(5). https://doi.org/10.1016/j.molcel.2020.12.047

Wallace, H. A. C., Marques-Kranc, F., Richardson, M., Luna-Crespo, F., Sharpe, J. A., Hughes, J., Wood, W. G., Higgs, D. R., & Smith, A. J. H. (2007). Manipulating the Mouse Genome to Engineer Precise Functional Syntenic Replacements with Human Sequence. Cell, 128(1). https://doi.org/10.1016/j.cell.2006.11.044

Wang, X., Cairns, M. J., & Yan, J. (2019). Super-enhancers in transcriptional regulation and genome organization. Nucleic Acids Research, 47(22), 11481–11496. https://doi.org/10.1093/NAR/GKZ1038

Whyte, W. A., Orlando, D. A., Hnisz, D., Abraham, B. J., Lin, C. Y., Kagey, M. H., Rahl, P. B., Lee, T. I., & Young, R. A. (2013). Master transcription factors and mediator establish super-enhancers at key cell identity genes. Cell, 153(2). https://doi.org/10.1016/j.cell.2013.03.035

Yamagata, K., Nakayamada, S., & Tanaka, Y. (2020). Critical roles of super-enhancers in the pathogenesis of autoimmune diseases. Inflammation and Regeneration, 40(1), 1–9. https://doi.org/10.1186/S41232-020-00124-9/TABLES/2

