## Supplementary figures for "Super-enhancers require a combination of classical enhancers and novel facilitator elements to drive high levels of gene expression"

### Supplementary material

A)

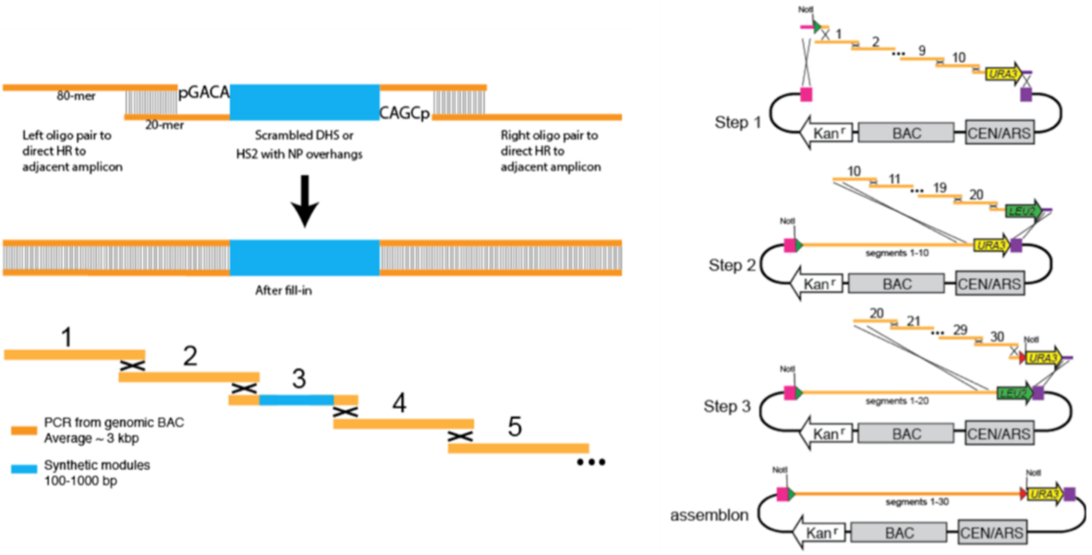


B)

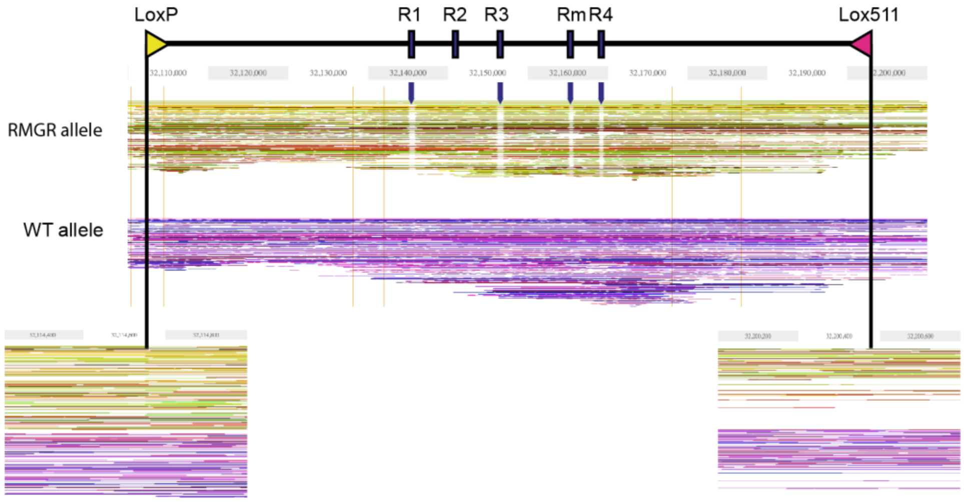


*Supplementary figure 1.*
*A) Short segments of DNA are generated by PCR from a target backbone or ordered as synthetic constructs with matching overhangs to allow for homologous recombination. Homologous recombination is performed in yeast with blocks of up to 10 segments in a single step. Alternating selection for the URA3 and LEU2 genes is used to perform subsequent assembly steps. The resulting BAC contains the assembled locus, modules necessary for replication in both yeast and E. Coli (BAC / CEN/ARS, respectively) and selectable markers including the kanamycin resistance gene (Kanr).*
*B) Top: schematic of the RMGR region with enhancer elements and RMGR lox exchange sites annotated. Middle: phased linked-reads for the R2-only RMGR and WT alleles, visualised with proprietary Loupe software from 10X Genomics. Enhancer element deletions are clearly visible by sequencing dropout in the RMGR allele, as indicated by arrows. Bottom: junctions where lox sites remain in the R2-only allele are seen as mapped read ends in the RMGR allele, but not in the WT allele where there are no lox sites.*

A)

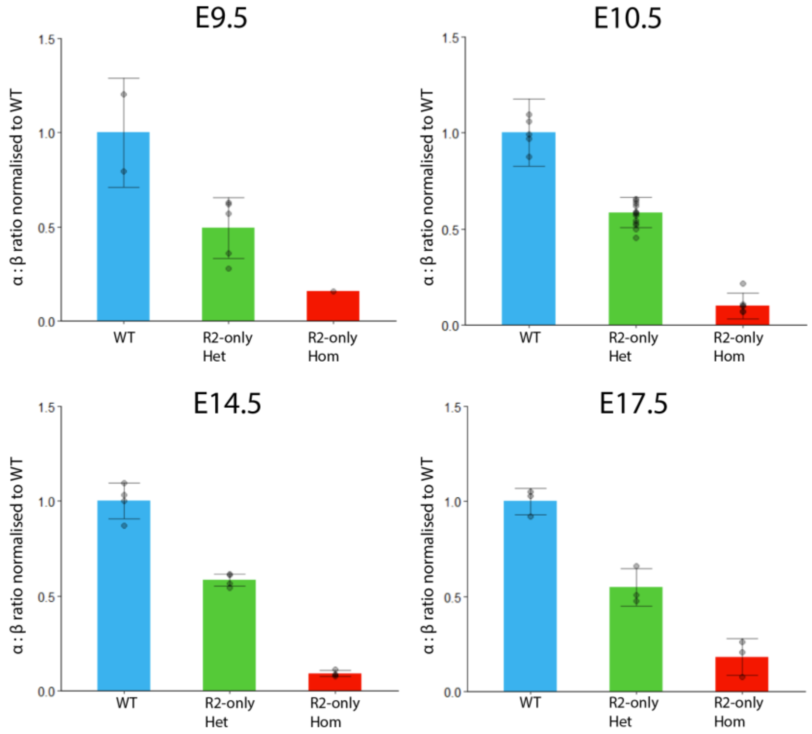


B)

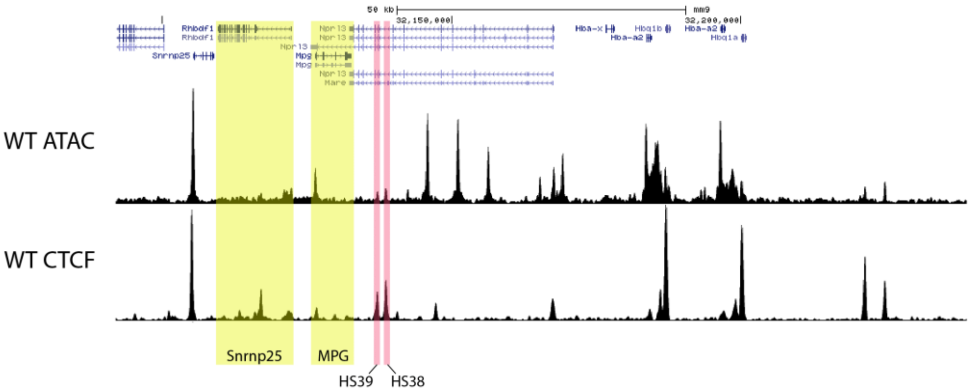


*Supplementary figure 2*
*A) RT-qPCR comparing α-globin expression in foetal liver erythroid cells from WT, R2-only heterozygous and R2-only homozygous E9.5. E10.5, E14.5 and E17.5 littermates. Expression normalized to β-globin and displayed as a proportion of WT expression. Dots = biological replicates; error bars = SE.*
*B) Top: ATAC-seq from WT E12.5 embryos. Bottom: CTCF ChIP-seq from WT E12.5 embryos. HS39 and HS38 CTCF (5' sub-TAD boundary) sites highlighted. Snrnp25 and MPG genes highlighted upstream of CTCF sites.*

A)

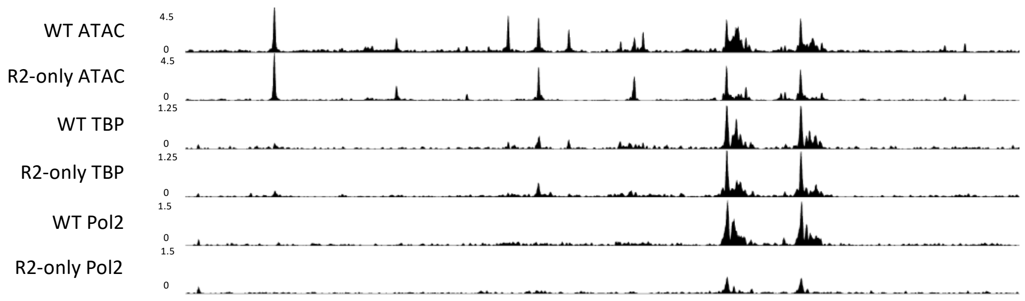


B)

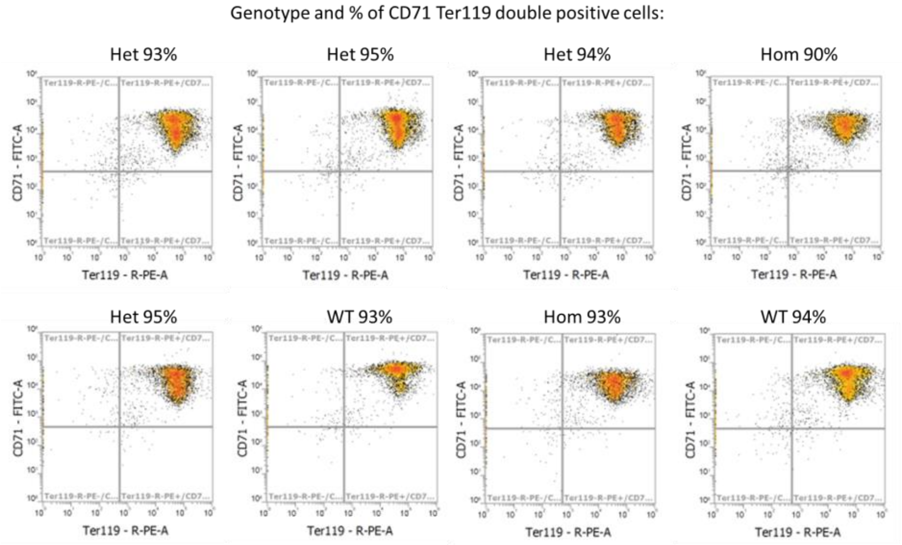


*Supplementary figure 3*
*A) ATAC-seq in WT (n=3) and R2-only homozygous (n=3) foetal liver erythroid cells; TBP ChIPmentation in WT (n=2) and R2-only homozygous (n=2) foetal liver erythroid cells; Pol2 ChIPmentation in WT (n=2) and R2-only homozygous (n=2) foetal liver erythroid cells. Tracks = merged biological replicates. (Coordinates = chr11: 32,090,000-32,235,000).*
*B) FACS from E12.5 foetal liver harvest, with antibodies against CD71 and Ter119. Percentage of CD71+/Ter119+ cells ~equivalent in WT, R2-only heterozygous and R2-only homozygous embryos.*

A)

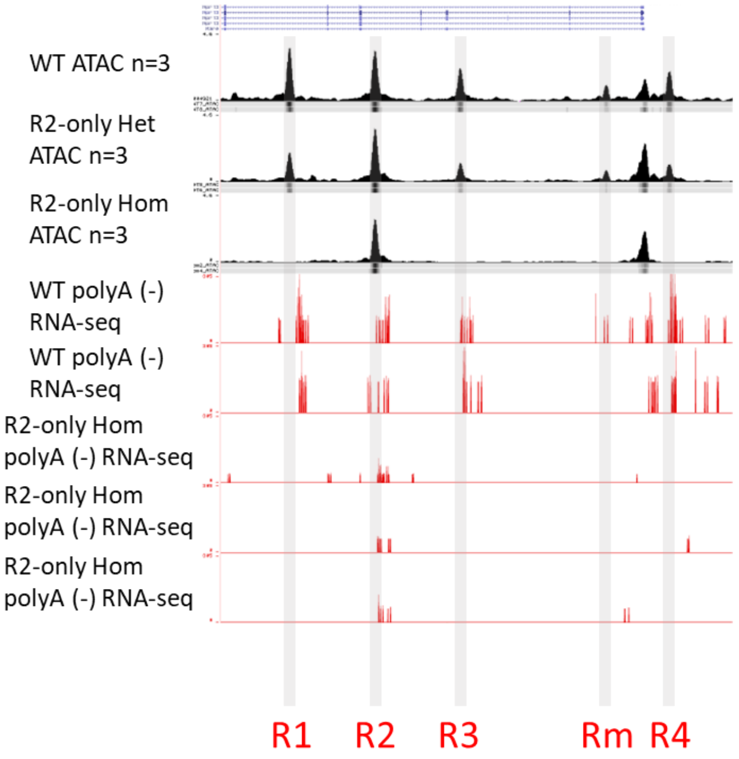


*Supplementary figure 4*
*A) ATAC-seq in WT (n=3), R2-only heterozygous (n=3) and R2-only homozygous (n=3) foetal liver erythroid cells; Poly A minus RNA-seq tracks from WT (n=2) and R2-only homozygous (n=3) foetal livers.*

*Supplementary table 1*
*Segregation of embryo genotypes following R2-only heterozygous X heterozygous crosses*

| Stage | Hom | Het | WT |
|---|---|---|---|
| **E9.5** | 1 | 5 | 2 |
| **E10.5** | 2 | 12 | 6 |
| **E12.5** | 5 | 7 | 5 |
| **E14.5** | 4 | 6 | 3 |
| **E17.5** | 3 | 7 | 6 |
| **Cumulative** | **15** | **37** | **22** |
| **Cumulative %** | **20** | **50** | **30** |

*Supplementary table 2*
*Guide RNA sequences used to delete each regulatory element.*

| gRNA | Sequence |
|---|---|
| R3 5′ | GGACTGAGGGAATTTTGTACA |
| R3 3′ | GGACTGGCAGAAAGCTATGT |
| Rm 5′ | GGACCAGCGTAGTCTAACTCC |
| Rm 3′ | GGACTCAACCACATGACTCA |
| R4 5′ | GGTGCATCACCTCAACTCCCA |
| R4 3′ | GGCTAGCTGAATAATTTGCGG |
| R4 HDR | GGATCCCAGGTTAGTTAGTCC |

*Supplementary table 3 Primer sequences used to genotype each regulatory element deletion*

| Target | Sequence (F/R) |
|---|---|
| R3 boundary PCR | TGTGAAATCACCAGAATTACAGG/ TGTGGGTCAGGCCTCTTAGAGGG |
| Rm boundary PCR | TAGGGAAAGGTTATGTGAACTGC/ TCAGGCTGGGCACTGGCTCTGCC |
| R4 boundary PCR | TTTGCCAGCAACCTTCACAGGAG/ ATTTTAAAGGCATTGCAGAGCCG |
| R2-only mutant PCR | AGCTCTGAAATGGACGTGGT/ GAGCCGTCCTGTGTATGTGA |
| R2-only WT PCR | GGGCACAGCAAAAGAGGAAA/ AATGGTCCTTGTCCTCCCAG |

*Supplementary table 4*
*Taqman assay IDs for RT-qPCR gene expression analysis*

| Target | Assay ID |
|---|---|
| Hba-a1/a2 | Custom (IDT) |
| Hba-x | Mm00439255_m1 |
| Hbb-bh1 | Mm00433932_g1 |
| Hbb-bh2 | Mm01273444_g1 |
| Hbb-bt/bs/b1/b2 | Custom (IDT) |
| Hbb-y | Mm00433936_g1 |
| Hbq1a | Mm00731011_s1 |
| Hbq1b | Mm02747875_s1 |
| Il9r | Mm00434313_m1 |
| Mpg | Mm00447872_m1 |
| Nprl3 | Mm01193449_m1 |
| Rhbdf1 | Mm00711711_m1 |
| Snrnp25 | Mm00547218_m1 |

*Supplementary table 5*
*Antibodies used in ChIPmentation analyses.*

| Antibody | ID |
|---|---|
| H3K27ac | Abcam ab4729 |
| H3K4me1 | Abcam ab8895 |
| H3K4me3 | Abcam 8580 |
| Gata1 | Abcam ab11852 |
| Nf-e2 | Santa Cruz sc-22827 |
| Med1 | Bethyl IHC-00149 |
| Brd4 | Bethyl IHC-00396 |
| Rad21 | Abcam Ab992 |
| CTCF | Cambridge Bioscience 61312 |
| PolII | Santa Cruz sc-899 |
